# Enhancing CDK4/6 inhibitor therapy for medulloblastoma using nanoparticle delivery and scRNA-seq-guided combination with sapanisertib

**DOI:** 10.1101/2021.06.09.447757

**Authors:** Chaemin Lim, Taylor Dismuke, Daniel Malawsky, Jacob D. Ramsey, Duhyeong Hwang, Virginia L. Godfrey, Alexander V. Kabanov, Timothy R. Gershon, Marina Sokolsky-Papkov

## Abstract

CDK4/6 inhibitors hold promise for brain tumor treatment, but efficacy has been limited by recurrence in both preclinical models and clinical trials. To address recurrence, we tested a nanoparticle formulation of the CDK4/6 inhibitor palbociclib (POx-palbo) in mice genetically-engineered to develop SHH-driven medulloblastoma. We then analyzed medulloblastomas in mice receiving palbociclib treatment, and compared the efficacy of combining palbociclib with specific inhibitors suggested by our analysis. POx-Palbo showed reduced toxicity compared to conventional palbociclib, was tolerable in parenteral administration, improved CNS pharmacokinetics, and extended survival of mice with medulloblastoma. Recurrence, however, remained problematic as fractions of tumor cells proliferated during therapy. ScRNA-seq identified a gene expression pattern unique to proliferating medulloblastoma cells in POx-Palbo-treated mice, marked by up-regulation of the glutamate transporter *Slc1a2* and down-regulation of diverse ribosomal genes. Reduced mTORC1 signaling, suggested by ribosomal suppression in POx-Palbo-treated tumors was confirmed by decreased 4EBP1 phosphorylation (p4EBP1). Further reducing mTORC1 activity by combining POx-Palbo with the mTORC1 inhibitor sapanisertib produced mutually enhancing effects, with increased suppression of both pRB and p4EBP1, and prolonged mouse survival compared to either agent alone. In contrast, targeting cell cycle progression by combining POx-Palbo with the SHH-pathway inhibitor vismodegib, or with the replication-targeting agents gemcitabine or etoposide, failed to enhance efficacy. Our data show the potential of nanoparticle formulation and scRNA-seq analysis of resistance to improve brain tumor treatment, and identify POx-palbo plus sapanisertib as effective combinatorial therapy for SHH medulloblastoma. This combination may be appropriate for testing in patients with recurrence, who need new options.

## INTRODUCTION

Patients with medulloblastoma, the most common malignant brain tumor in children, need new therapies. The current standard treatment for medulloblastoma with surgery, craniospinal radiation (xRT) and chemotherapy cures ~80% of patients, but causes disabling untoward effects, including neurocognitive impairment, hearing loss, endocrine dysfunction and secondary malignancies. Survivors remain at risk of recurrence after treatment, and recurrent medulloblastoma is presently incurable. Therapies that specifically target the biology of the tumor and spare normal tissues may improve the efficacy of medulloblastoma therapy and decrease long-term toxicities.

CDK4/6 inhibitors may be ideal targeted agents for medulloblastoma (*1*, *2*). While medulloblastomas are a heterogeneous group of tumors with 4 subgroups, all 4 groups of medulloblastomas typically disable the RB tumor suppressor through CDK4/6-mediated RB phosphorylation (pRB) and *RB* mutations are rare in every subgroup (*3*–*5*). The CDK4/6 inhibitor palbociclib (Ibrance, Pfizer, Inc.) effectively blocks RB phosphorylation and arrests cells with intact RB in the G_1_ phase of the cell cycle (*6*) and is FDA-approved for specific breast cancers, in combination with hormonal therapy (*7*, *8*).

Implementing palbociclib for medulloblastoma therapy may require both improving CNS pharmacokinetics (PK) and identifying mutually enhancing drug combinations. The brain penetration of palbociclib is limited (*9*), and dose-limiting toxicities restrict the potential for increasing systemic doses to improve CNS drug delivery. Additionally, alternative mechanisms can compensate for CDK4/6 activity, as cells proliferate effectively in *Cdk4/6*-deleted mouse embryos (*10*). These mechanisms may support palbociclib resistance in tumors, and combinations of drugs may be needed to block resistance mechanisms (*11*). Consistent with these obstacles, palbociclib efficacy has been limited by recurrence in mouse models of diffuse, intrinsic pontine glioma (*12*), and medulloblastoma (*1*, *2*). Similarly, palbociclib was ineffective as a monotherapy in Phase II clinical trials for recurrent glioblastoma patients with detectable RB expression (NCT01227434) (*13*). Dose-limiting toxicities may have contributed to poor clinical efficacy, as the tested doses were low relative to doses in preclinical studies and many patients required dose reductions due to neutropenia. At the doses that were tolerable for patients, no pharmacodynamic (PD) effect was noted and all patients recurred during therapy (*13*). Addressing the problems of CNS drug delivery and mechanisms of resistance may realize the potential of palbociclib for medulloblastoma, and more broadly for brain tumor therapy.

We have previously shown that polyoxazoline (POx) based amphiphilic block copolymer micelles can act as nanoparticle carriers of diverse small molecules for CNS delivery (*14*, *15*). POx delivery of the SHH inhibitor vismodegib increases brain and tumor drug exposure, reducing systemic toxicity and increasing efficacy (*16*). Based on these prior studies, we tested whether POx delivery improved palbociclib. We designed a novel POx block copolymer to effectively load palbociclib and compared the resulting formulation (POx-Palbo) to conventional palbociclib in the treatment of SHH-driven medulloblastomas that developed spontaneously in *Smo*-mutant mice, analyzing pharmacokinetics, pharmacodynamics, toxicity and efficacy. To address medulloblastoma recurrence during therapy, we investigated the cell cycle progression and single-cell gene expression profiles of proliferating tumor cells in POx-Palbo-treated medulloblastomas. We then tested combinations of POx-Palbo with specific agents suggested by our analyses. Our results show that POx delivery improved the CNS PK and efficacy of palbociclib, identify palbociclib-induced changes in gene expression in the tumors cells that proliferated during therapy, and show that combining POx-Palbo with the mTOR inhibitor sapanisertib dramatically increased efficacy compared to either agent alone.

## RESULTS

Limited efficacy of conventional palbociclib in medulloblastoma suggests the need for nanoparticle drug delivery to the CNS

We found that conventional palbociclib (Palbo-HCl) was ineffective in a genetic model of aggressive, refractory SHH medulloblastoma, indicating the need for optimization. We generated mice with SHH medulloblastoma by breeding *hGFAP-Cre* mice that express Cre reombinase in CNS stem cells during development with *SmoM2* mice that express a Cre-conditional oncogenic allele of *Smo*. The resulting *G-Smo* mice developed medulloblastoma with 100% penetrance by P10 and, untreated, died of progressive tumors by P20, as in prior studies (*17*, *18*). We determined the maximum tolerated doses (MTD) for healthy WT mouse pups treated daily with oral Palbo-HCl starting at P10, or parenteral Palbo-HCl, administered IP daily from P10-P14, then every other day (Fig. 1a). We then treated *G-Smo* mice starting at P10 with similar regimens of Palbo-HCl. Compared to saline-injected controls, Palbo-HCl produced no statistically significant increase in the event-free survival of *G-Smo* mice when administered orally at either the MTD of 100 mg/kg/day or 50% of the MTD or when administered parenterally at the MTD of 10 mg/kg/day. Considering the poor efficacy of Palbo-HCl in our model, we determined whether nanoparticle drug delivery might improve drug performance.

**Fig. 1.**
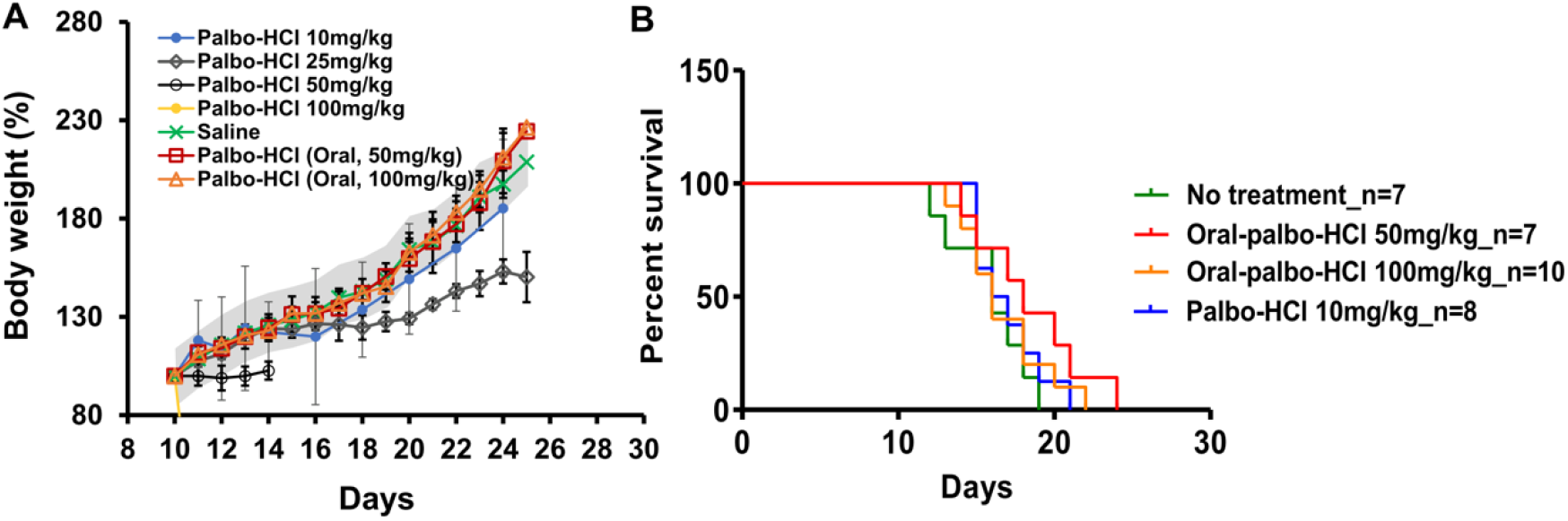
**(A)** Maximal Tolerated Dose (MTD) of Palbo-HCl following oral or systemic administration. Palbo-HCl was administered daily at the indicated doses starting at P10, and mice were weighed each day. The gray range indicates the mean weights SEM of uninjected littermate controls. The MTD was defined as the highest dose that did not reduce weight gain by >15%. **(B)** Kaplan-Meir curve of *G-Smo* mice treated with the indicated regimens. Oral Palbo-HCl was dosed daily starting on postnatal day 10. Palbo-HCl IP was.

To develop a nanoparticle formulation, we attempted to load palbociclib into the poly(2-oxazoline polymer, that we previously used to formulate vismodegib (POx-A) (*16*). This hydrophilic, non-charged A-B-A type copolymer, (P[MeOx_34_-*b*-BuOx_20_-*b*-MeOx_35_] (M_n_ = 8.6 kg/mol)) showed very low loading capacity **(Fig. 2A)**. Noting that palbociclib is a pyridopyrimidine that forms hydrophobic as well as hydrogen bonds due to its secondary piperazine nitrogen (pK_a_ 7.4) and pyridine nitrogen (pK_a_ 3.9) groups which act as hydrogen donor/acceptors, we designed new POx block co-polymers predicted to have stronger polymer-drug interactions. We synthesized three different copolymers with carboxylic groups on the side chains and varying hydrophobicity: POx-B: hydrophilic A-B-A type copolymer (P[MeOx_38_-*b*-PpaOx_27_-*b*-MeOx_38_], M_n_ = 10.6 kg/mol), POx-C: an A-B-C copolymer, (P[MeOx_33_-*b*-BuOx_21_-*b*-PpaOx_29_], M_n_ = 9.9 kg/mol) with hydrophilic methyl and hydrophobic butyl groups, and the most hydrophobic POx-D: A-B-C copolymer with hydrophobic ethyl and butyl blocks, (P[P[EtOx_34_-*b*-BuOx_21_-*b*-PpaOX_31_], M_n_ = 10.7 kg/mol) **(fig. S1)**. We then assessed the ability of each polymer to encapsulate palbociclib using the thin film method.

**Fig. 2.**
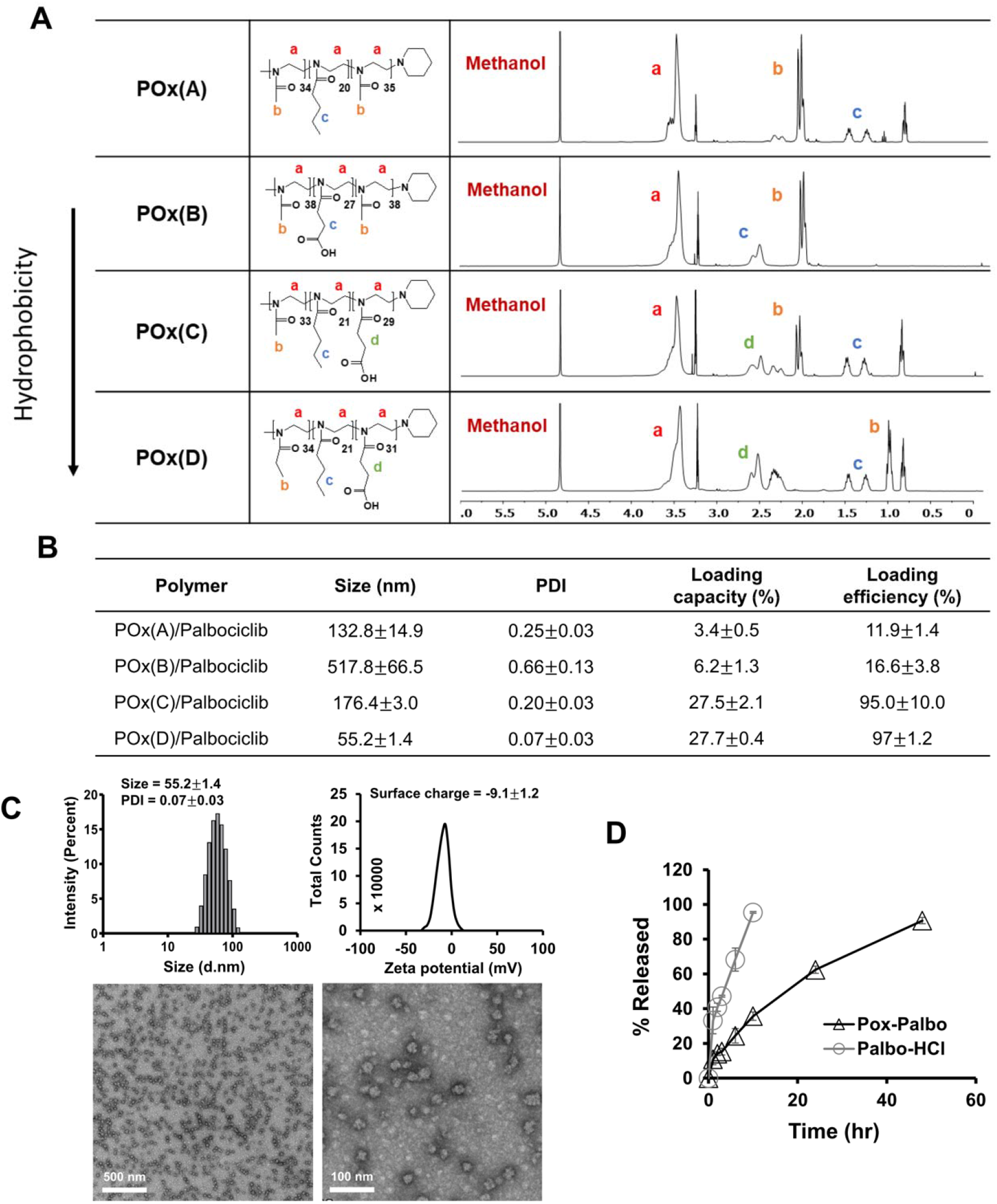
Specific block co-polymers improve the loading of palbociclib into polyoxazoline micelle nanoparticles. **(A)** Strucutres of poly(2-oxazoline) triblock copolymers. The structures were cofirmed by ^1^H NMR spectrum (in MeOD) by indenfying NMR peaks. **(B)** Particle size, size distribution and loading parameters (LC%, LE%) of palbociclib-loaded micelles prepared from indicated polymers. Loading capacity (%) = M_drug_ / (M_drug_ + M_excipient_) × 100 %, Loading efficiency (%) = M_drug_/ (M_drug added_) × 100 % (n = 3 ± SD). **(C)** Particle size distribution (z-average, Dz, by DLS), Zeta potential and morphology (by TEM, scale bar = 500 nm (left), 10 nm (right)). **(D)** Palbociclib release profile from POx-Palbo incubated in 10% fetal bovine serum (FBS) solution at at 37 over time. Palbo-HCl was used as penetration control.

Introducing carboxylic groups alone, as in POx-B, was not sufficient to improve drug loading. Adding hydrophobic blocks, as in POx-C and POx-D, effectively improved drug loading to ~28% LC. The further increase in hydrophobicity in POx-D compared to POx-C, reduced the particles size and size distribution (**Fig. 2B**). The POx-D-palbociclib micelles were small, spherical particles with mean size of 55nm and narrow size distribution (PDI < 0.1) as determined by dynamic light scattering (DLS) and transmission electron microscopy (TEM; **Fig. 2C).** We selected the POx-D formulation as optimal based on loading capacity and particle size and designated it as POx-Palbo for further studies. *In vitro* release studies of POx-Palbo showed a sustained release profile without burst release, with approximately 30% of palbociclib released in 6 hours and 90% released within 48 hours **(Fig. 2D)**.

### POx micelle formulation reduced systemic toxicity after parenteral administration

We compared the tolerability of POx-Palbo versus Palbo-HCl in MTD studies. We injected healthy, non-tumor bearing pups intraperitoneally with escalating doses of POx-Palbo daily on postnatal days 10-14 (P10-14) and then every other day until P26, as in the parenteral Palbo-HCl regimen and analyzed toxicity, defined as 15% less weight gain compared to saline-treated littermate controls. The MTD for POx-Palbo was 50 mg/kg (**Fig. 3A**) markedly higher than the 10 mg/kg MTD for Palbo-HCl. POx-Palbo therefore showed superior tolerability in parenteral administration.

**Fig. 3.**
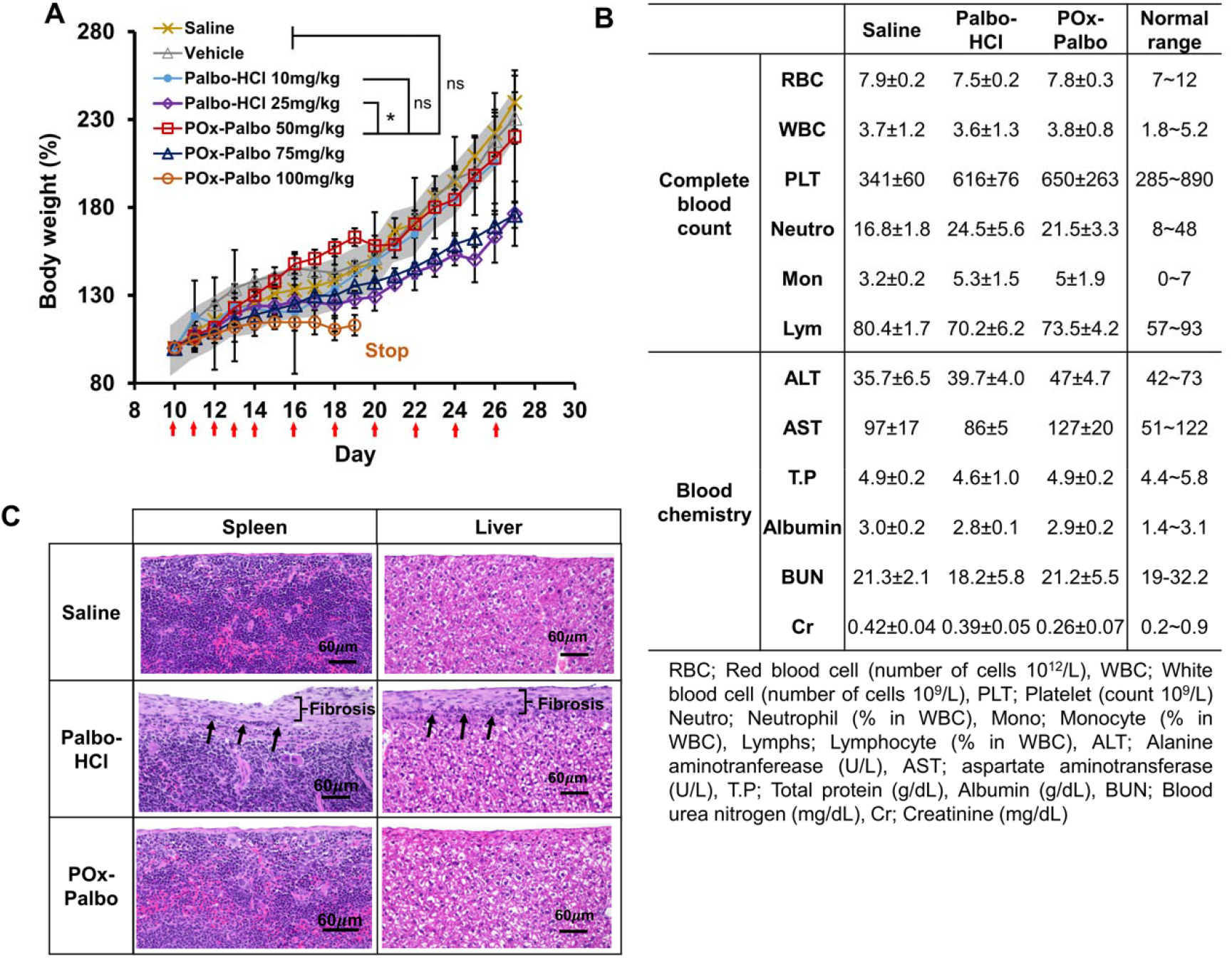
POx-Palbo reduces toxicity. (A) The weights of mice treated with the indicated formulations over time. The gray range indicates the mean weights of ± SEM of littermate controls (*p < 0.05). (B) Complete blood count and clinical chemistry parameters and (C) Hemotoxylin & eosin (H&E) staining of C57BL/6 mice treated with saline, Palbo-HCl, or POx-Palbo (equivalent palbociclib 25 mg/kg). The square bracket and arrows indicate the fibrosis and ongoing inflammatory cells respectively. Mice were treated IP daily on postnatal days 10-14 (P10-14) and then every other day until P24, and blood samples and tissues were collected 24h after the last injection (P25) (n = 3 ± SD).

To identify toxicities, we treated mice with 25 mg/kg of Palbo-HCl, the highest dose that did not produce fatal toxicity, or with an equivalent dose of POx-Palbo, using the regimen of daily administration on postnatal days 10-14 (P10-14) and then every other day until P24. We found no significant changes in complete blood counts (CBC) or serum levels of liver enzymes, total protein, albumin, blood urea nitrogen and creatinine compared to saline-treated controls (**Fig. 3B**). Histologic examination of the heart, thymus, lung, liver, spleen, and kidney demonstrated fibrosis of the peritoneal organs in all Palbo-HCl treated mice, that was not seen in POx-Palbo-treated mice **(Fig. 3C)**. This fibrosis indicated peritoneal inflammation that likely explains the poor tolerability of IP Palbo-HCl. While IP tolerability is not relevant to clinical implementation, the absence of fibrosis with POx-Palbo treatment provides important evidence that the drug remained in the nanoparticle carrier until after systemic uptake.

### POx micelles improved palbociclib delivery to the brain and brain tumors

We compared the pharmacokinetics of Palbo-HCl and POx-Palbo in the blood, forebrain and medulloblastomas of *G-Smo* mice. For these comparisons, we administered each formulation at a constant palbociclib dose of 25 mg/kg, selected as the highest IP dose of Palbo-HCl that did not produce fatal toxicity. We incorporated tritiated palbociclib into POx-Palbo and Palbo-HCl formulations administered these tritium-labeled agents to groups of replicate *G-Smo* mice on P10. We then harvested the mice at successive time points after administration and collected plasma and tissue samples. We measured palbociclib concentrations by scintillation counting and analyzed the results using Phoenix Modeling software.

Palbo-HCl showed shorter times to peak drug concentrations (T_max_) in plasma and all tissues sampled, and higher volume of distribution (8.45 mL compared to 3.07 mL for POx-Palbo (p-value <0.05). While C_max_ in plasma was comparable between the formulations, in tumors the C_max_ of POx-Palbo was 75% higher compared to Palbo-HCl (5.45 μg/g compared to 3.28 μg/g) and the POx-Palbo AUC was nearly two-fold greater (p-value <0.05) (**Fig. 4 A** **and** **B**). Palbo-HCl produced higher concentrations in the kidneys consistent with renal clearance, while POx-Palbo reached higher peak concentrations in the liver concentrations **(table S1)**, consistent with typical nanoparticle clearance in the hepatobiliary system (*19*). Compartment modelling highlighted the differences between the free drug and micellar formulation. Palbo-HCl fit a two-compartment model while the POx-Palbo micelles better fit a one compartment model, suggesting reduced exposure to unintended tissues.

**Fig. 4.**
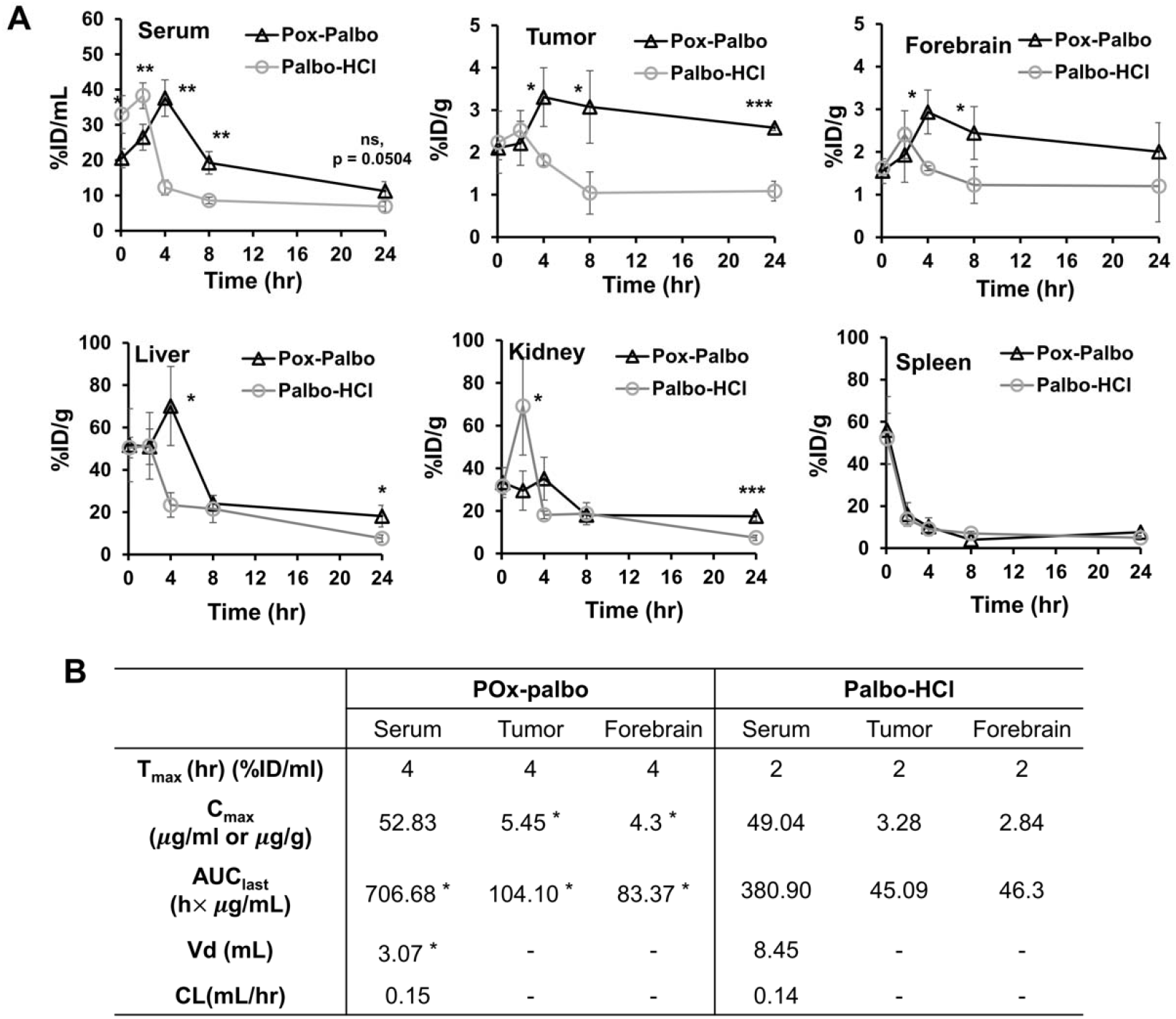
Pharmacokinetic profile of POx-Palbo and Palbo-HCl in medulloblastoma-bearing *G-Smo* mice. **(A)** Plots of palbociclib concentration in serum and major organs (Tumor, forebrain, liver, kidney and spleen over time). **(B)** PK parametes of given formulations in serum, tumor and forebrain (n = 3 ± SD). Statistical analysis was performed with t-tests, *p < 0.05, **p < 0.01, ***p < 0.001.

### POx-Palbo shows a significant anti-tumor effect, consistently limited by recurrence

To determine if the improved PK of POx-Palbo correlated with improved efficacy *in vivo*, we compared the event free survival times of *G-Smo* mice treated with regimens of 50 mg/kg IP daily (the MTD), 25 mg/kg IP daily (50% of MTD), or 25 mg/kg IP daily for 7 days, followed by 12.5 mg/kg thereafter (**Table s2**). The lower dose POx-Palbo regimens were used to determine if less intense, less toxic doses would differently effect animal survival. All POx-Palbo regimens showed similar efficacy with increased animal survival compared to *G-Smo* treated saline as sham controls or treated with Palbo-HCl at the MTD of 10 mg/kg IP daily (**Fig. 5 A**). POx-Palbo-treated mice consistently showed smaller tumors day P15 (**Fig. 5B**), but eventually progressed during therapy, indicating that efficacy waned over time. To gain insight into the mechanisms of resistance and recurrence, we analyzed POx-Palbo pharmacodynamics, and determined whether the pharmacodynamics changed over the course of therapy.

**Fig. 5.**
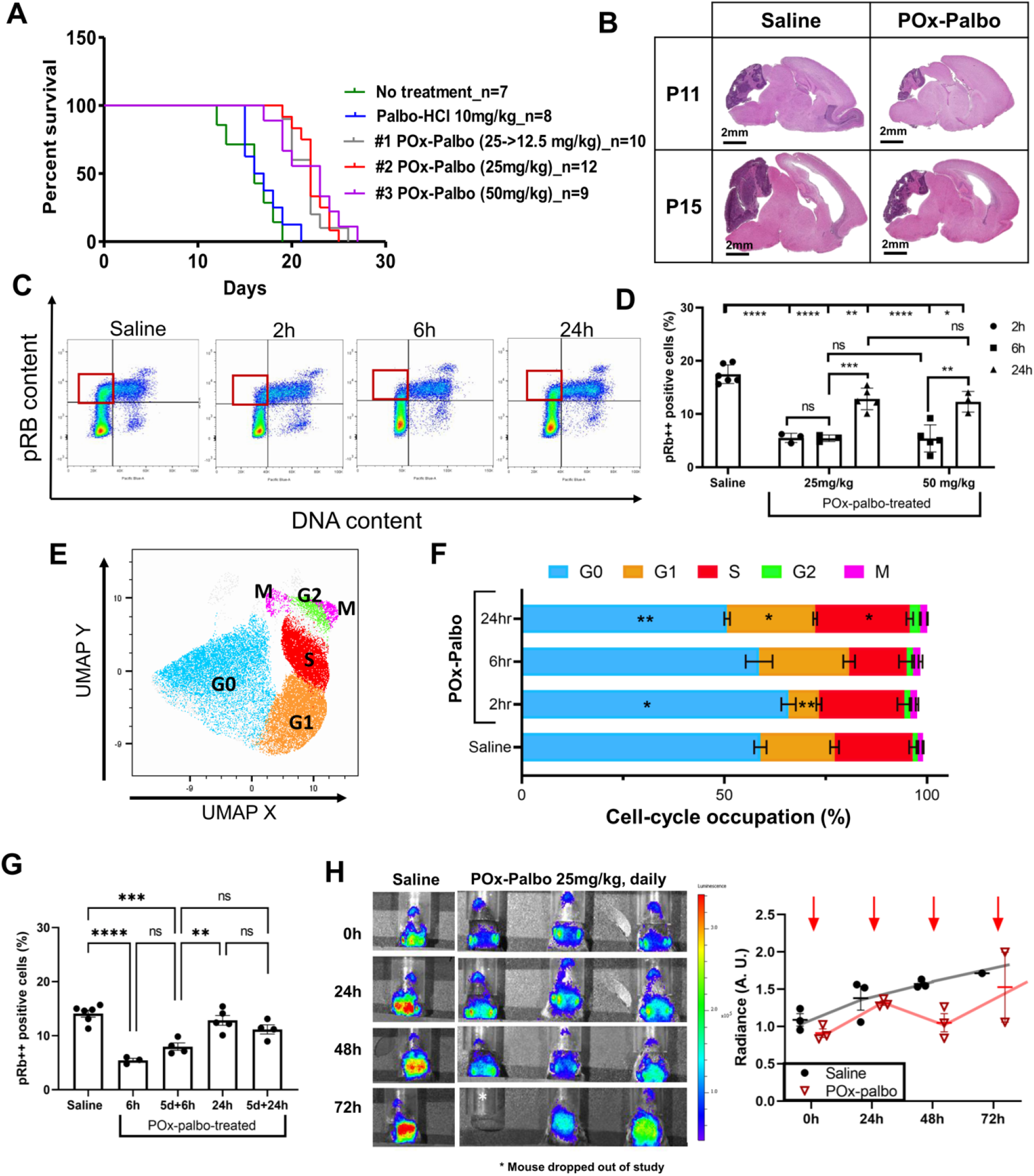
Pharmacodynamic analysis of POx-Palbo in medulloblastoma- bearing *G-Smo* mice. **(A)** Kaplan-meir curve of POx-Palbo treatments (#1-3) and palbo-HCl in *G-Smo* mice. **(B)** H&E of POx-Palbo treated brains at 24 hours (P11) and 5 daily treatments (P15) compared to saline controls. **(C)** Two-dimensional flow cytometry analysis of DNA content (x-axis) and pRB content (y- axis) in POx-Palbo treated (2-24h) *G-Smo* mice. **(D)** Quantification of highly phosphorylated cells in G1 (Red square) in POx-Palbo-treated mice. Statistics performed with One-way ANOVA. **(E)** UMAP qualitative map of tumor cells dissociated from *G-Smo* mice. **(F)** Cell-cycle occupation analysis of 25 mg/kg of POx-Palbo treated (2-24h) *G-Smo* mice. Unpaired t-test of each cell-cycle phase compared to saline control. **(G)** Quantification of highly-phosphorylated cells in G1 in 25 mg/kg of POx-Palbo treated mice after 5 days. Statistical analysis was performed with One-way ANOVA. (H) Longitdunial dynamic tracing of POx-Palbo in luciferase-tumor bearing mice. Quantified in the right panel. *p < 0.05, **p < 0.01, ***p < 0.001, ****p < 0.0001.

### POx-Palbo pharmacodynamics remain stable as efficacy decreases

We determined pharmacodynamics after *in vivo* POx-Palbo treatment by quantifying RB phosphorylation and cell cycle progression. We administered POx-Palbo IP to *G-Smo* mice, then harvested tumors at defined intervals after drug administration, injecting EdU IP 30 minutes before harvest to label cells at S-phase. We then dissociated the tumors, fixed and stained the cells for pRB, EdU and DNA content, and used flow cytometry to quantify cells at each phase of the cell cycle. We idnetified G_0_ as pRB- cells with 2N DNA content, G_1_ as pRB+/EdU- cells with 2N DNA content, S-phase as EdU+/pRB+ cells, and G_2_/M as EdU- cells with 4N DNA content (**Supplementary Fig. 2**). The fraction of pRB+ cells with 2N DNA content, which included cells in G_1_ and early S-phases, was markedly reduced in POx-Palbo-treated tumors at both 2 and 6 hours after administration (**Fig. 5 C** **and** **D**), demonstrating effective inhibition CDK4/6, as expected. By 24 hours, pRB+ cells with 2N DNA were significantly lower compared to controls, but significantly higher compared to 6 hours after administration, indicating waning CDK4/6 inhibition(**Fig. 5D**). Increasing the dose of POx-Palbo from 25 mg/kg to 50 mg/kg did not increase pRB suppression (**Fig. 5D**), indicating that the dose of 25 mg/kg saturated the capacity of the system to respond.

In contrast to pRB, cell cycle progression decreased in the hours after POx-Palbo administration, but more than fully resumed by 24 hours (**Fig. 5 E,F)**. POx-Palbo-treated tumors showed significantly reduced cells at G1 at 2 hours after injection, and reduced cells at S-phase at 6 hours after injection, indicating that fewer cells proceeded from G_0_ to S-phase (**Fig. 5F**). However, by 24 hours, POx-Palbo-treated tumors showed significantly increased G_1_, S and G_2_/M cells, indicating a compensatory increase in cell cycling (**Fig. 5F**). While cell cycle progression resumed more rapidly than RB phosphorylation, the suppression on both processes decreased more rapidly than tumor drug concentration and the waning pharmacodynamics therefore could not be explained by pharmacokinetics alone.

To determine if medulloblastoma sensitivity to palbociclib was affected by repeated exposure, we treated mice with daily POx-Palbo 25 mg/kg IP for 5 days and then harvested replicate tumors at successive intervals after the fifth dose and analyzed pharmacodynamics using our flow cytometry methods. We found that medulloblastomas treated for 5 days with daily POx-Palbo showed similar temporal patterns of pRB suppression after the fifth dose, indicating that the magnitude and duration of CDK4/6 inhibition were not changed by repeated exposure over this period **(Fig. 5G)**.

While repeated doses of POx-Palbo continued to inhibit CDK46, longitudinal studies of tumor growth during therapy showed diminishing effects over the same time period. We measured tumor growth non-invasively *in vivo* by breeding *Gli-luc* transgenic mice that report SHH activity through a GLI-sensitive luciferase reporter (*20*), into the *G-Smo* model as in our prior studies (*21*). The resulting *G-Smo^Gli-luc^* mice developed medulloblastomas detectable by bioluminescence imaging (BLI). BLI signal in untreated *G-Smo^Gli-luc^* mice increased progressively from P10-P14 (**Fig. 5H**, **control**) indicating growth of SHH-activated tumors. Daily treatment of replicate *G-Smo^Gli-luc^* mice with POx-Palbo 25 mg/kg IP produced an initial decrease in mean BLI, indicating tumor suppression. By day 4, however, the tumor suppressive effect of POx-Palbo diminished as BLI signal began increasing while on daily therapy **(Fig. 5H)**. Tumor growth therefore resumed during the period of stable pharmacodynamics, indicating that the recurrence did not require loss of CDK4/6 inhibition or resistance to pRB suppression, but rather a dissociation between pRB suppression and growth suppression. To define the processes that allow medulloblastoma cells to proliferate during ongoing palbociclib treatment, we compared the tanscriptomic profiles of proliferative medulloblasted cells in untreated *G-Smo* tumors versus tumors in *G-Smo* mice on POx-Palbo therapy, using scRNA-seq.

### scRNA-seq studies show that proliferative tumor cells in palbociclib-treated tumors down-regulate ribosomal genes

We used Drop-seq (*22*) methods as in our prior studies (*21*, *23*) to compare tumors isolated from three P15 *G-Smo* mice treated with 25 mg/kg POx-Palbo daily IP for 5 days to tumors from five age-matched, untreated *G-Smo* mice. Briefly, tumors were harvested, dissociated and individual cells were paired with bar-coded, oligo dT-coated beads using microfluidic methods, and bar-coded cDNA was synthesized on the beads. We then prepared and sequenced amplified libraries from the pooled cDNA and used cell-specific bar codes to identify the transcriptomes of individual cells in the resulting sequence data. We filtered cells identified by bead-specific bar codes to address unintentional cell-cell multiplexing and premature cell lysis (*24*, *25*). 1530 out of 4959 cells from POx-Palbo treated tumors, and 8699 out of 16489 cells from control tumors met criteria and were included in the analysis. To compare the two groups at similar sequencing depths, we randomly down-sampled the control transcript counts to 20% of the original depth (*26*).

We performed an integrated analysis of scRNA-seq data from POx-Palbo treated and control tumors using PCA as in our prior studies (*21*), and generated a 2-dimensional UMAP projection in which individual cells were placed according to their similarities, forming clusters of mutually similar cells **(Fig. 6A)**. As in our prior medulloblastoma scRNA-seq studies, the UMAP showed both discrete clusters and a set of clusters with shared borders. We identified each cluster using markers of proliferation and CGNP lineage to identify tumor cells, and using markers of different types of stromal cells expected in the brain, as validated in our prior medulloblastoma scRNA-seq studies (*21*, *23*). These markers showed that each discrete cluster comprised a different type of stromal cell including astrocytes, oligodendrocytes, myeloid cells, endothelial cells and fibroblasts (**Fig. 6 A** **and** **B**; **Table 1)**. Marker analysis identified the 6 clusters with shared borders as medulloblastoma cells in a range of differentiation states with opposite proliferative and differentiated poles. Proliferation markers *Mki67* and *Pcna*, and ghe SHH-driven transcripiton factor *Gli1* identified the cells of clusters 0,1,3 and 5 as proliferative; similarly intermediate differentiation markers *NeuroD1* and *Cntn2*, and late differentiation markers *Grin2b* and *Gria2*, placed the cells of clusters 2 and 4 in successive states of differentiation **(Fig. 6 C,D).**

**Fig. 6.**
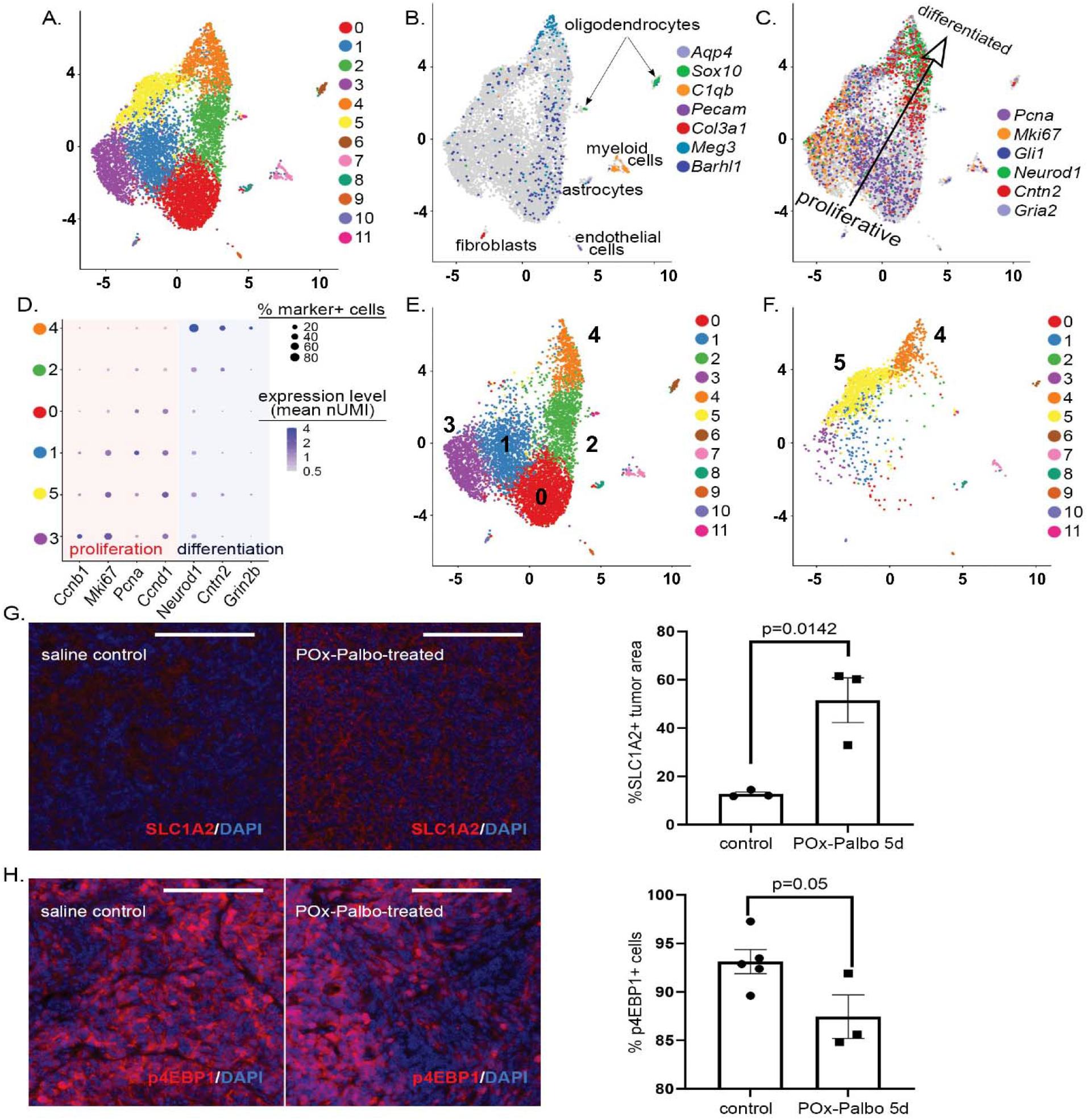
scRNA-seq shows key differences in the composition of POx-Palbo-treated tumors, including a population of proliferative medulloblastoma cells not found in control tumors, marked by up-regulation of *Slc1a2* and down-regulation of mTORC1 activity. **(A)** UMAP projection of cells from POx-Palbo-treated and control tumors, grouped by transcriptomic similarities into color-coded clusters. **(B)** Expression of indicated markers color-coded onto the UMAP projection from **(A)**, with stromal cluster identities indicated. **(C)** Expression of proliferation and differentiation markers is color-coded onto the UMAP projection from **(A)**. The arrow indicates the overall direction of progression from the proliferative to the differentiated pole. **(D)** Dot Plot shows the magnitude and frequency of the expression of indicated proliferation and differentiation markers in the indicated tumor cell clusters. **(E,F)** UMAP from **(A)**, disaggregated by condition to show **(E)** cells from control tumors and **(F)** cells from POx-Palbo-treated tumors. **(G)** Representative sections of control and POx-Palbo treated *G-Smo* tumors, immunostained for SLC1A2, with quantification. **(H)** Representative sections of control and POx-Palbo-treated *G-Smo* tumors, immunostained for p4EBP1, with quantification. The p values in **(G,H)** determined by Student’s t-test. Scale bars = 100 μm.

**Table 1.**
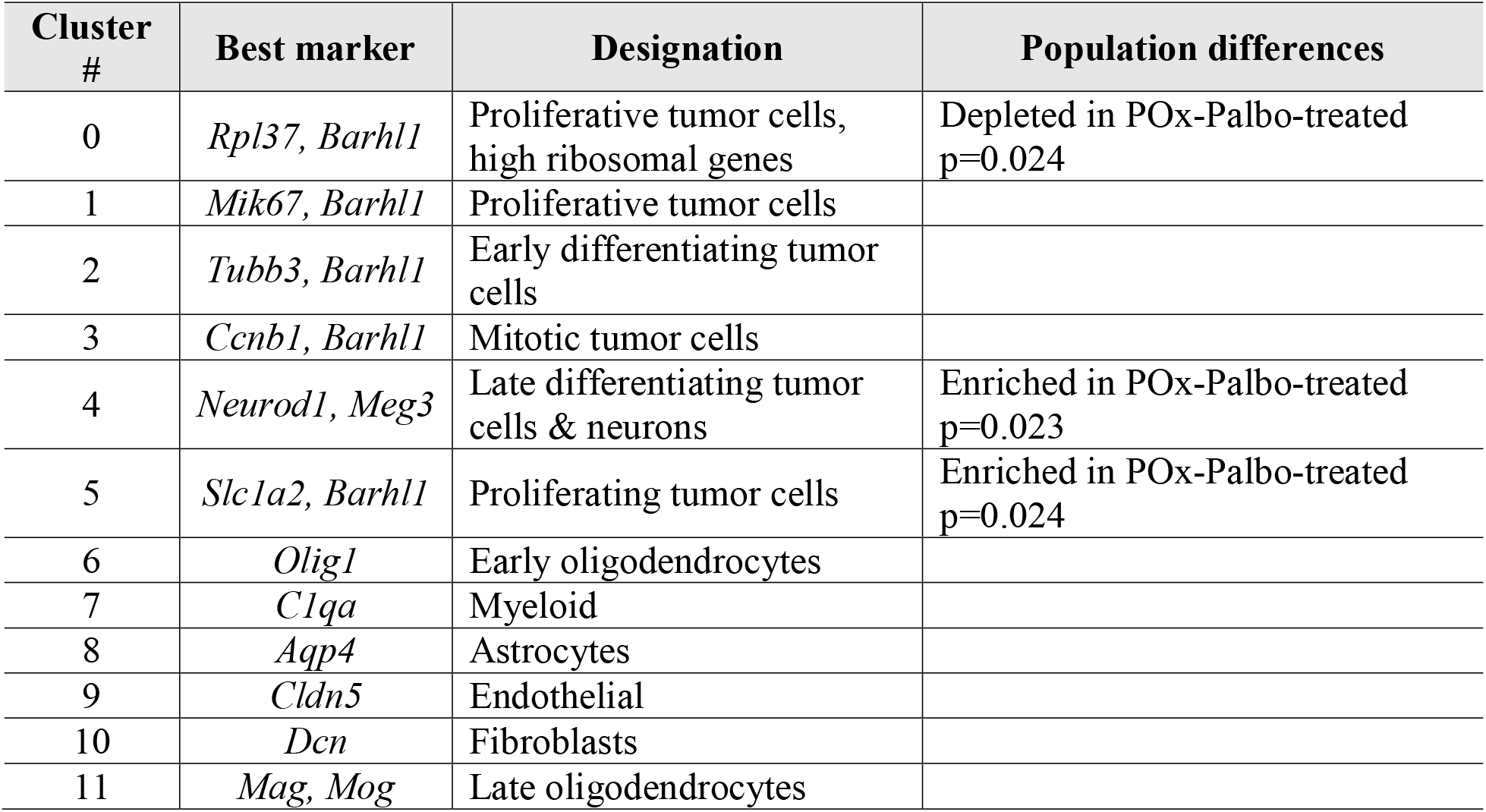
Cluster identities, markers, and population differences by condition.

Disaggregating the data by treatment group showed that medulloblastoma cells were not evenly distributed in the tumor cell clusters **(Fig. 6 E,F)**. Palbociclib-treated tumors contained significantly larger fractions of cells in cluster 4 (p=0.023), the most differentiated cluster. Proliferative medulloblastoma cells were also differentially distributed across clusters. Cluster 5 specifically comprised palbociclib-treated cells (p=0.024, Wilcoxon rank sum test) and cluster 0 comprised predominantly control cells (p=0.024), while clusters 1,3 did not show statistically significant between treatment groups. POx-Palbo-treated tumors thus contained both more differentiated tumor cells, consistent with drug-induced growth-suppression, and a distinct population of proliferating cells with a transcriptomic profile that was different from proliferating cells in control tumors.

We considered that the gene expression patterns that distinguished cluster 5 cells from the other proliferative cells may include both potentially growth-suppressive mechanisms that failed to block proliferation and resistance mechanisms that allowed proliferation to continue on therapy. To identify these patterns, we determined which genes were differentially expressed in cluster 5 cells, compared to the cells of clusters 0,1 and 3, then filtered to include only the set of genes that was expressed by a 2-fold or greater proportion of cluster 5 cells, or the set expressed by a 2-fold or greater proportion of cells of clusters 0, 1, and 3 **(Data file S1)**. We subjected both sets to Gene Ontology (GO) enrichment analysis to identify biological processes discernably affected.

GO analysis of the set of genes enriched in clusters 0, 1, and 3 identified translation as the most strongly increased process (p=1.2 × 10^−39^), with diverse biosynthetic processes also increased. GO analysis of the set of genes enriched in cluster 5 cells identified less specific terms including “positive regulation of biologic process” and “positive regulation of metabolic process” as the most significantly increased (p=6.5 × 10^−9^ and 6.8 × 10^−9^ respectively), with chromatin organization also significantly increased. Consistent with increased translation in clusters 0, 1, and 3, we noted increased expression of diverse ribosomal genes and *Eef1b2* **(Data file S1)**. Consistent with metabolic and chromatin alterations induced by palbociclib, we noted increased expression of the glutamate transporter *Slc1a2* and the chromatin modifier *Smarca4* in cluster 5 cells **(Data file S1)**. We confirmed increased protein expression of SLC1A2 (aka GLT1) in replicate POx-Palbo treated and control tumors using immunohistochemistry (IHC) **(Fig. 6 G)**.

Signaling through mTORC1 regulates ribosomal gene transcription as part of a general regulation of translation (*27*), and we therefore analyzed whether mTORC1 activity was reduced in cells that remained proliferative during chronic palbociclib treatment. Inhibition of mTORC1 has been shown to increase SLC1A2 expression in other cell types (*28*), which also suggested reduced mTORC1 activity in POx-Palbo treated medulloblastomas. To investigate mTORC1 activation, we analyzed phosphorylated 4EBP1 (p4EBP1) as a marker of mTORC1 activity. We compared p4EBP1 in replicate *G-Smo* mice treated for 5 days with POx-Palbo versus untreated *G-Smo* controls, using IHC. We found significantly fewer p4EBP1+ cells in POx-Palbo-treated tumors, indicating reduced mTORC1 activation, consistent with decrease in ribosomal genes in the scRNA-seq data **(Fig. 6H)**.

These data show reduced mTORC1 activity and increased SLC1A2 protein expression in palbociclib-treated medulloblastomas, validating our scRNA-seq data. These correlations do not show whether reduced mTORC1 activity or increased SLC1A2 are direct effects of the drug or secondary effects of pRB suppression, or whether the effects are mechanisms of resistance that allows continued proliferation. We hypothesized, however, that whether reduced mTORC1 activity was growth-suppressive or growth-enabling, the mTORC1 activity in POx-Palbo-treated tumors might already be at the lower limit of tolerability for proliferating cells, resulting in increased sensitivity to small molecule mTORC1 inhibitors. We therefore tested whether combining POx-Palbo with the mTORC1 inhibitor sapanisertib, which, like palbociclib shows medulloblastoma efficacy limited by recurrence (*29*), would produce an efficacy greater than either agent alone.

### Testing combination therapy with POx-Palbo plus multiple agents validated POx-(Palbo+Sapanisertib) and failed to show benefits of other combinations

In parallel with the palbociclib-sapanisertib combination, we tested the potential of agents targeting SHH signaling or DNA replication to limit resistance when combined with palbociclib. We theorized that vismodegib, which inhibits the SHH receptor component SMO, would combine favorably with palbociclib by targeting SHH-driven proliferation at two distinct points, SMO and CDK4/6. We further theorized that replication-targeted agents gemcitabine or etoposide would enhance palbociclib efficacy by disrupting tumor cells that progress to S-phase despite CDK4/6 inhibition, and reciprocally that the compensatory increase in cycling cells 24 hours after palbociclib would increase gemcitabine and etoposide sensitivity. We tested these hypotheses by developing regimens for each agent, then treating cohorts of *G-Smo* mice with each agent, either individually, or in combination with POx-Palbo.

To facilitate administration of vismodegib, etoposide, sapanisertib, which were not water-soluble, we developed POx-based micellar formulations. The use of POx-formulations was supported by our prior findings that vismodegib formulated in polyoxazoline nanoparticles (POx-Vismo) was more effective than conventional vismodegib when administered to *G-Smo* mice (*16*) and that etoposide loaded into POx micelles (POx-Etop) was less toxic and more effective than free drug in mouse lung cancer models (*30*, *31*). We characterized these formulations by size, size distribution and drug loading **(fig. S3A).** For gemcitabine studies, we used a conventional, rather than POx formulation, as gemcitabine is water soluble and did not load well into polyoxazoline micelles. The dose of each single drug was determined based on MTD evaluations *in vivo* in healthy mice without tumors. These studies identified 5 mg/kg gemcitabine every three days as the MTD **(fig. S3B)**. The MTD for POx-Etop was 5 mg/kg every 5 days **(fig. S3C)**, however at this dose mice showed marginally acceptable weight gain, and we therefore used 2.5 mg/kg (50% of the MTD) for tumor treatment. The MTD for formulation of sapanisertib loaded into POx micelles (POx-Sapanisertib) was 0.2 mg/kg daily **(fig. S3D)**; as with etoposide we used 0.1 mg/kg dosing due to marginal weight gain at the MTD. For combinational studies, we co-encapsulated the pairs of drugs in single nanoparticle formulations. The drugs ratios were varied to evaluate the effects of drug ratios on treatment outcomes (**fig. S4**).

To combine vismodegib and palbociclib, we administered POx-(Palbo+Vismo) starting at P10 using 3 different treatment schedules, including reducing POx-Palbo to 12.5 mg/kg after 5 doses and increasing Vismo dose to 100 mg/kg **(Table 2**; **table s3)**. We tested these different regimens to identify the most effective, tolerable combination. None of tested regimens, however, were superior to either POx-Palbo or Pox-Vismo as single agents (**Fig. 7A**). Next, we tested 3 different schedules of combined gemcitabine and POx-Palbo **(Table 2**; **table s4)**; all failed to improve survival compared to Pox-Palbo alone **(Fig. 7B)**.

**Table 2.**
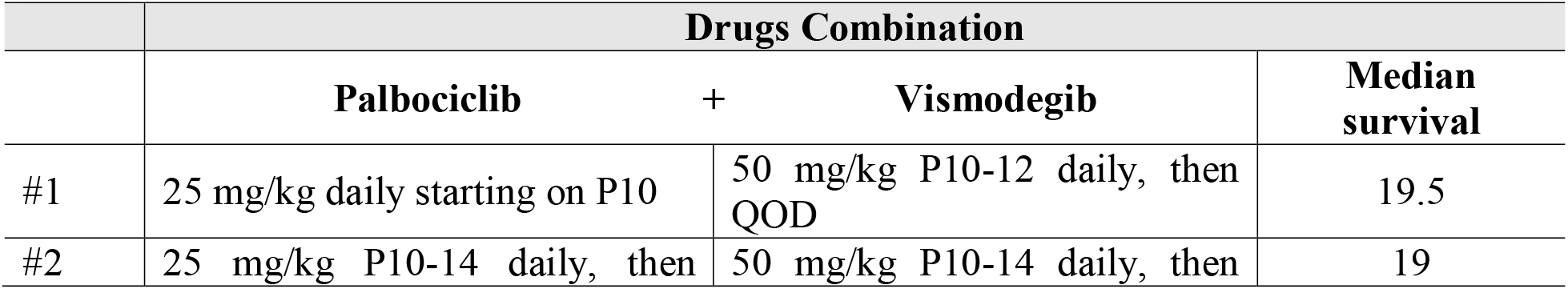
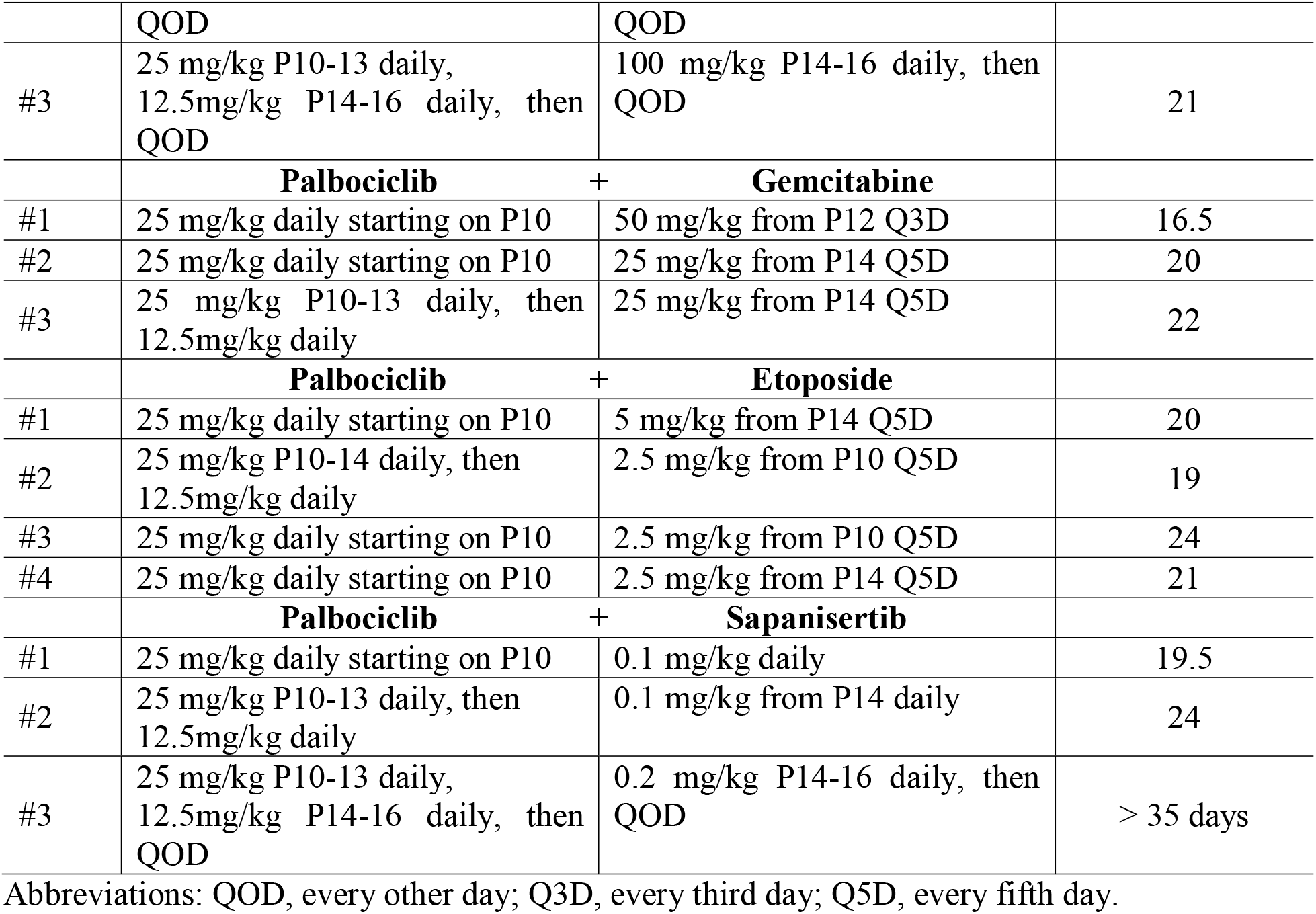
Regimens used in *in vivo* testing

**Fig. 7.**
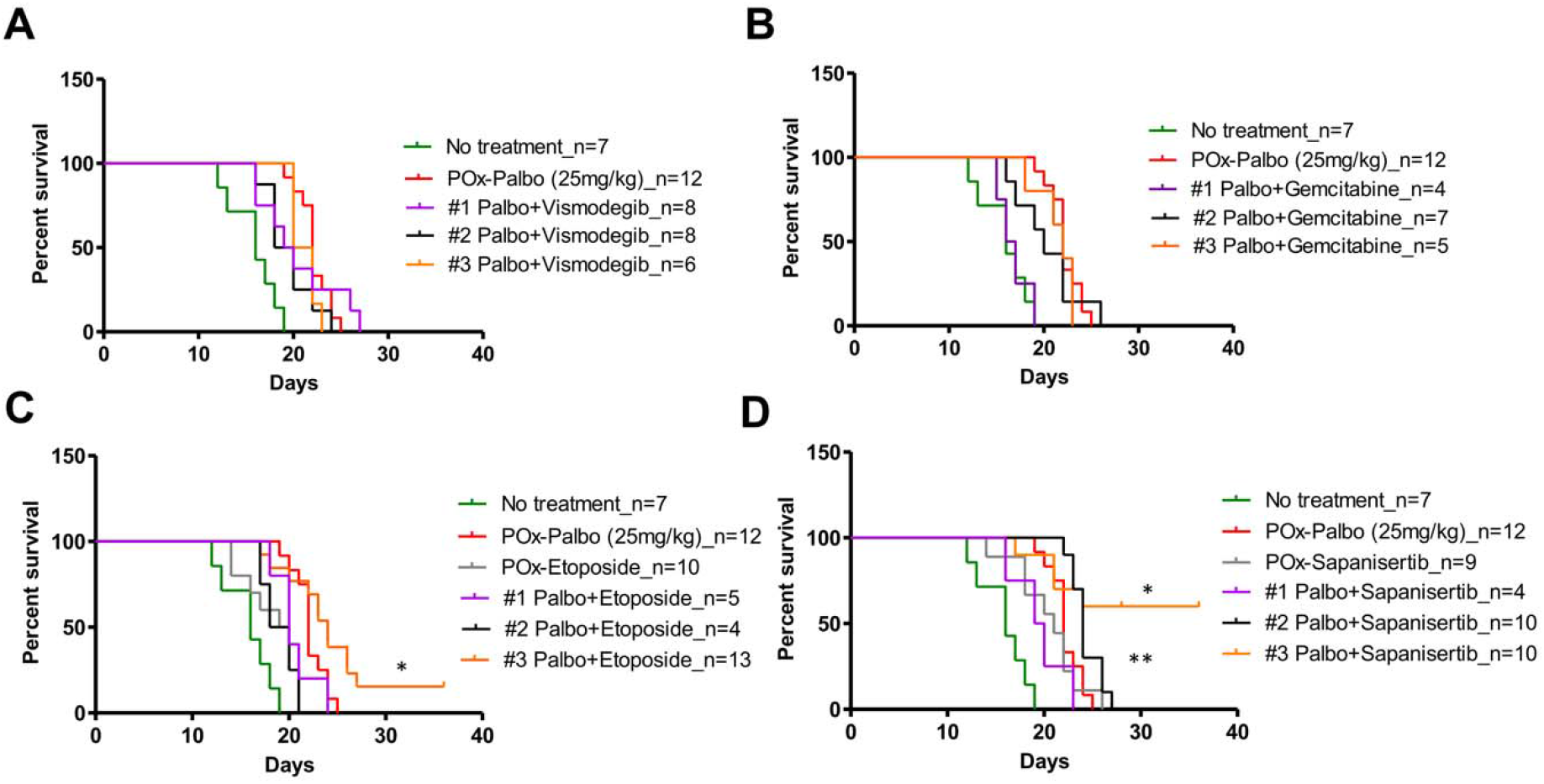
Kaplan-Meier survival curves for *G-Smo* mice treated with (A) POx-(Palbo+Vismo) (B) POx-Palbo+Gemcitabine (C) POx-(Palbo+Etop), (D) POx-(Palbo+Sapanisertib) in the indicated regimens. The p values are determined by comparing each combination to POx-Palbo single-agent therapy, using the Log-rank (Mantel-Cox) Test. *p < 0.05, **p < 0.01.

We tested multiple regimens with varied doses of etoposide and palbociclib **(Table 2**; **table s5)**. Treatment of *G-Smo* mice with single agent POx-Etop 2.5 mg/kg IP every 5 days improved survival compared to sham treatment **(p = 0.03; Fig. 7C)**. The regimen of POx-(Palbo+Etop) (25 mg/kg palbociclib/2.5 mg/kg etoposide) every 5 days starting on P10 with POx-Palbo 25 mg/kg daily between combined doses improved mouse survival relative to both POx-Palbo alone (p=0.047) and POx-Etop alone (p=0.001). Recurrence during therapy, however, remained a consistent limitation **(Fig. 7C)**.

We tested multiple regimens of sapanisterib and palbociclib **(Table 2**; **table s6)**. Single agent POx-Sapanisertib dosed at 0.1 mg/kg IP to *G-Smo* mice significantly improved survival compared to sham treatment **(Fig. 7D)**. Mice on combined regimen 1 were ill-appearing, with reduced spontaneous movement, and we therefore reduced the dose of palbociclib to 12.5 mg/kg after day 4 of treatment in regimens 2 and 3, and further reduced the frequency of administration in regimen 3 to every other day, increasing the sapanisertib component to 0.2 mg/kg to maintain the same sapanistertib dose over time. Regimens 2 and 3 both showed improved survival compared to POx-Palbo alone. Regimen 3 was most effective, superior to both POx-Palbo alone (p=0.011) and POx-Sapanisertib alone (p=0.006), and reduced recurrence during therapy from 100% to 40% **(Fig. 7D)**. Combining palbociclib with mTORC1 inhibitor sapanisertib was therefore markedly more effective in reducing recurrence during treatment than all other combinations tested.

### Palbociclib and sapanisertib show enhanced pharmacodynamics

Pharmacodynamic studies showed combining palbociclib and sapanisertib enhanced the mechanistic effects of each agent. We compared the temporal patterns of pRB suppression and p4EBP1 suppression after administration of POx-Palbo, POx-Sapanisertib or POx-(Palbo+Sapanisertib) using flow cytometry and immunohistochemistry **(Fig. 8)**. 6 hours after administration, POx-Palbo, POx-Sapanisertib and POx-(Palbo+Sapanisertib) significantly and similarly decreased pRB; however, pRB suppression by POx-(Palbo+Sapanisertib) was markedly more durable, remaining significantly reduced compared to controls 24 hours after administration **(Fig. 8A–C)**.

**Fig. 8.**
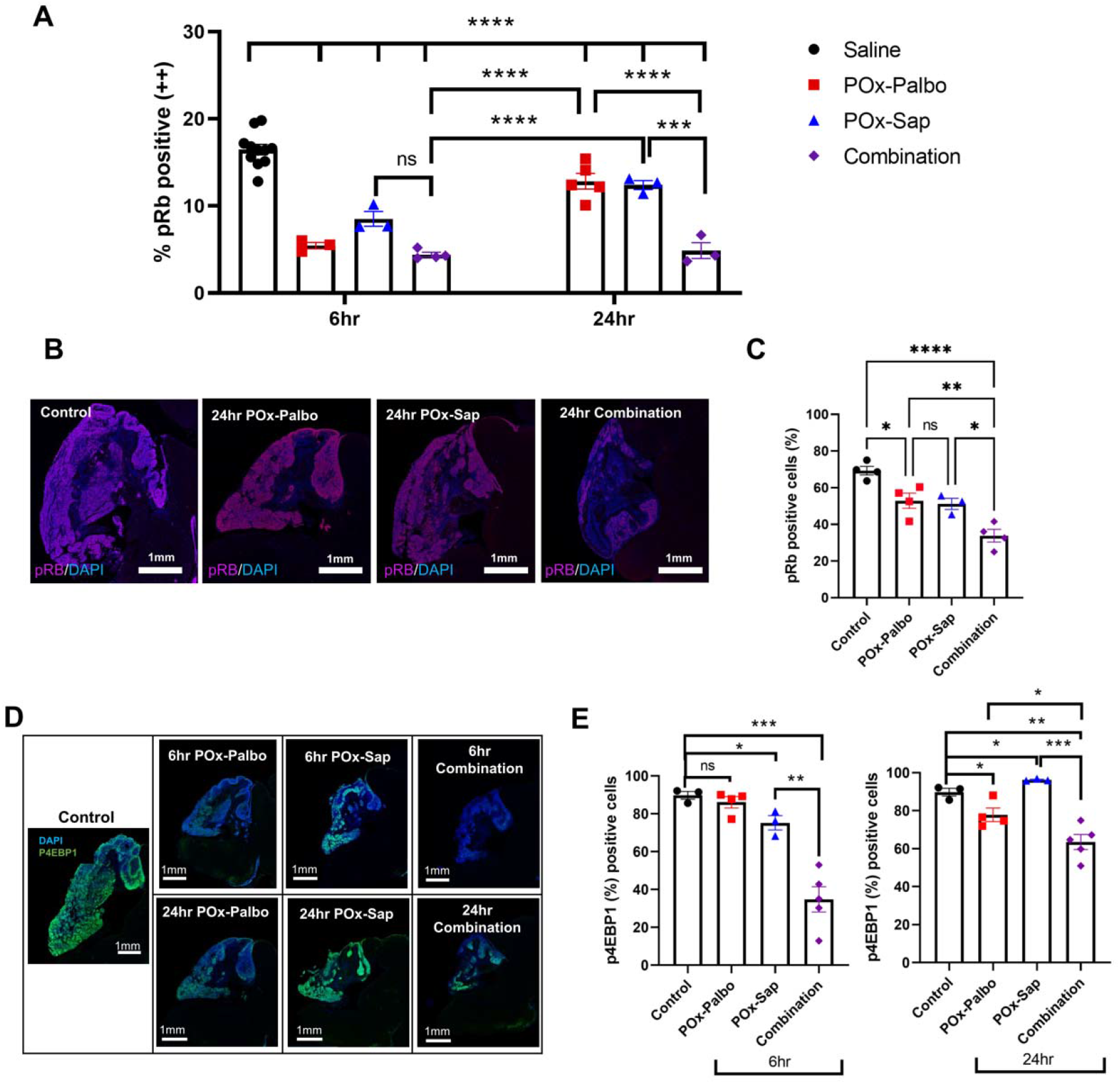
Pharmacodynamic analysis of POx-(Palbo+Sapanisertib) in medulloblastoma-bearing *G-Smo* mice. **(A)** Quantification of flow cytometry analysis of pRB+ G_1_ cells in POx-Palbo (red), POx-sapanisertib (blue), and POx-(Palbo+Sapanisertib) (purple) in *G-Smo* mice at 6 and 24 hours after adminstration. The p values were determined by one-way ANOVA. **(B)** Representative pRB immunofluorescence in *G-Smo* medulloblastomas 24 hours after the indicated treatment. **(C)** Quantification of pRB immunofluorescence in replicate mice, as in **(B)**. **(D)** Representative p4EBP1 immunofluorescence in *G-Smo* medulloblastomas 6 or 24 hours after the indicated treatment. **(E)** Quantification of phospho-4EBP1 immunofluorescence in replicate mice, as in **(D)**. The p values were determined by Student’s t-test.

Both POx-Sapanisertib and POx-(Palbo+Sapanisertib) significantly reduced p4EBP1 6 hours after administration, but p4EBP1 suppression by POx-(Palbo+Sapanisertib) was more durable and persisted over 24 hours **(Fig. 8 D,E)**. The enhanced pharmacodynamic effects of palbociclib and sapanisertib, delivered as POx-(Palbo+Sapanisertib), support the increased efficacy of the combination that was predicted by the scRNA-seq data and demonstrated by the survival studies.

## DISCUSSION

CDK4/6 inhibitor therapy for brain tumors has been complicated by suboptimal pharmacokinetics and the development of recurrence during therapy. We developed a polyoxazoline micelle-based nanoparticle formulation of the CDK4/6 inhibitor palbociclib, POx-Palbo, which allowed parenteral administration and improved CNS pharmacokinetics. Parenteral POx-Palbo improved mouse survival in the *G-Smo* model of aggressive, refractory SHH-driven medulloblastoma. In contrast Palbo-HCl failed to improve survival of G-Smo mice with either oral or parenteral Palbo-HCl administration. Recurrent disease, however, consistently limited POx-Palbo efficacy. While pharmacodynamic analysis of G-Smo mice treated for 5 days showed that palbociclib continued to inhibit CDK4/6 and to suppress pRB, cell cycle progression resumed by 24 hours after each dose, despite reduced pRB and tumor growth resumed within 5 days of the start of treatment. Analysis of medulloblastomas in POx-palbo-treated mice using scRNA-seq showed specific transcriptional changes in the tumor cells that remained proliferative during palbociclib treatment, including up-regulation of the glutamate transporter *Slc1a2* and suppression of ribosomal genes indicating of mTORC1 inhibition. Further decreasing mTORC1 activity by adding the mTORC1 inhibitor sapanisertib to the palbociclib regimen markedly increased anti-tumor efficacy. Similarly enhanced efficacy was not observed on combining palbociclib with vismodegib or gemcitabine; etoposide enhanced the efficacy of palbociclib, but less effectively than sapanisertib despite several dose optimization attempts. These data show the potential for nanoparticle technology to optimize CNS drug delivery, and for transcriptomic analysis with single cell resolution to identify processes that when targeted therapeutically can reduce recurrence.

The reduced ribosomal gene expression pattern that we observed in *G-Smo* tumors treated with POx-Palbo *in vivo* matches the patterns of differentially suppressed transcripts previously reported in medulloblastoma cell lines with CDK6 deletion and in *Cdk6*-deleted medulloblastomas that form in *Math1-Cre/SmoM2/Cdk6^−/-^* mice (*32*). Our data show that this transcriptomic change correlates with decreased mTORC1 activation, demonstrated by reduced p4EBP1. The reduced ribosomal function in *Cdk6*-deleted SHH medulloblastomas has been shown to sustain proliferation by inducing production of SMO-stimulating lipids (*32*). This mechanism may explain the failure of palbociclib to combine well with the SMO inhibitor vismodegib, as SMO-activating lipids induced by POx-Palbo may antagonize the inhibition of SMO by vismodegib. In contrast, by reducing mTORC1, which allows medulloblastoma cells to proliferate in the context of reduced CDK4/6 activity, palbociclib increased susceptibility to mTORC1 inhibitors. These data show that when mechanisms of resistance involve downregulation of fundamental biologic processes such as reduced mTORC1 activity, resistance may be blocked by amplifying rather than inhibiting the resistance mechanism, as in the addition of sapanisertib to palbociclib.

The combination of palbociclib and sapanisertib, suggested by our scRNA-seq and p4EBP1 studies, has been similarly reported to produce enhanced anti-tumor activity in a mouse model of intrahepatic cholangiocarcinoma (ICC) induced by somatic gene transfer of mutant Akt and Yap (*33*). In these ICC tumors, palbociclib alone produced a transient anti-tumor effect, followed by resistance mediated by increased CCND1, and addition of sapanisertib reduced CCND1 expression and potentiated cell cycle inhibition. In medulloblastomas, we found that palbociclib resistance did not involve increased *Ccnd1* expression. Sapanisertib, however, potentiated pRB suppression, as in ICC. In both *G-Smo* medulloblastomas and in ICC, single-agent therapy with palbociclib reduced p4EBP1, indicating mTORC1 inhibition, and also potentiated p4EBP1 suppression when combined with sapanisertib (*33*). A potential mechanism for the potentiating effect of palbociclib on sapanisertib-mediated mTORC1 inhibition is suggested by the suppression of CDK4-mediated activation of IRS2 and resulting inactivation of TSC2 (*11*). The mutually enhancing effects of palbociclib and sapanisertib in both SHH medulloblastoma and ICC show that this combination may be effective in diverse cancers driven by different oncogenic pathways in different tissues of origin.

## MATERIALS AND METHODS

### Materials

All materials for the synthesis of poly(2-oxazoline) block copolymers, methanol, ethanol, and sodium lactate were purchased from Sigma Aldrich (St. Louis, MO). Water and acetonitrile (HPLC grade) were purchased from Fisher Scientific Inc. (Fairlawn, NJ). All materials for the synthesis of poly(2-oxazoline) block copolymers including methyl trifluoromethanesulfonate (MeOTf), 2-methyl-2-oxazoline (MeOx), 2-ethyl-2-oxazoline (EtOx), 2-n-butyl-2-oxazoline (BuOx), and 2-methoxycarboxyethyl-2-oxazoline (MestOx) were dried by refluxing over calcium hydride (CaH_2_) under inert nitrogen gas and subsequently distilled prior to use. Palbociclib free base, palbociclib-HCl salt form, vismodegib, etoposide, and gemcitabine were purchased from LC Laboratories (Woburn, MA). Sapanisertib was purchased from Medkoo Biosciences (Morrisville, NC). [^3^H] Palbociclib was purchased from American Radiolabeled Chemicals (St. Louis, MO). Soluene-350 and Ultima Gold LLC scintillation cocktail were purchased from PerkinElmer Life and Analytical Sciences (Waltham, MA).

### Synthesis of POx block copolymers

Triblock copolymers, POx(A) (P[MeOx_34_-*b*-BuOx_20_-*b*-MeOx_35_] (M_n_ = 8.6 kg/mol)), POx(B) (P[MeOx_38_-*b*-PpaOx_27_-*b*-MeOx_38_] (M_n_ = 10.6 kg/mol)), POx(C) (P[MeOx_33_-*b*-BuOx_21_-*b*-PpaOx_29_] (M_n_ = 9.9 kg/mol)), and POx(D) (P[EtOx_34_-*b*-BuOx_21_-*b*-PpaOX_31_] (M_n_ = 10.7 kg/mol)) were synthesized by step-by-step (A-B-C) living cationic ring-opening polymerization (Scheme S1). Under dry and inert conditions, 1 equiv of MeOTf and pre-calculated equiv of corresponding block A monomer were dissolved in dry acetonitrile. The mixture was reacted in the microwave for 15min at 150 W and 130 °C. After cooling to room temperature, the monomer for block B was added and reacted again for another 15 min. The procedure was repeated with the monomer for block C and the polymerization was terminated by addition of 3 equiv of piperidine and incubating for 1hr at 50W and 40 °C. To remove the methyl ester group from P(MestOx), each polymer was dissolved in MeOH and mixed with 0.1N NaOH (1 eq to methyl ester group) at 90 °C for 3hr. The final solution was dialyzed (MWCO, 3500) against distilled water for 2 day and freeze-dried.

^1^H NMR spectrum was obtained using INOVA 400 and analyzed using MestReNova (11.0) software. The spectra were calibrated using the MeOD solvent signals (4.78 ppm). The number-average molecular weight was determined by ^1^H NMR, by calculating the ratio of the initiator and each repeating unit, using samples taken upon polymerization of each block and final block copolymer.

### Preparation of drug loaded POx micelles

Drug-loaded polymeric micelles were prepared by thin-film hydration method as previously described. Each polymer and drug stock solutions in methanol were mixed together at the pre-determined ratios, followed by complete evaporation of methanol under a stream of nitrogen gas. The well-dried thin films were subsequently rehydrated with normal saline and then incubated at room temperature to self-assembly into drug-loaded polymeric micelles. The resulting micelle solutions were centrifuged at 10,000 g for 3 min (Sorvall Legend Micro 21R Centrifuge, Thermo Scientific) to remove non-loaded drug. The concentration of drugs in micelles were analyzed by reversed-phase high-pressure liquid chromatography (Agilent Technologies 1200 series) with a Nucleosil C18, 5 μm column (L × I.D. 250 mm × 4.6 mm). Samples were diluted 20 times in mobile phase (mixture of acetonitrile/water, with 0.01% trifluoroacetic acid) and 10 *μ*L of the diluted sample was injected into the HPLC while the flow rate was 1.0 mL/min and column temperature was 40°C. The retention time of drugs, detection wavelength and detailed mobile phase were presented in **table S7**. The drug loading efficiency (LE) and loading capacity (LC) were calculated using following equations.

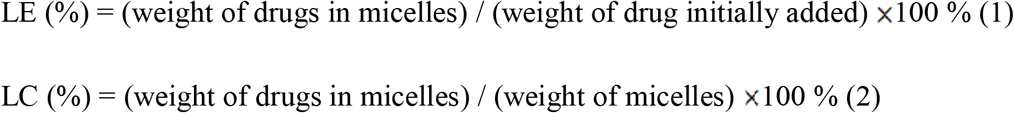

### Characterization of drug loaded POx micelles

The particle size, polydispersity index (PDI) and zeta potential of drug loaded POx micelles were measured by photon correlation spectroscopy using a Zetasizer Nano-ZS (Malvern Instruments, Worcestershire, UK). Before measurement, each micelle was diluted to yield 1 mg/mL final polymer concentration and the average values were calculated from three independent sample measurements. To observe the morphologies of POx micelles, one drop of micelle solution (diluted 100 times using distilled water) was placed on a copper grid/carbon film and stained with negative staining (1% uranyl acetate) prior to the TEM imaging.

The release profile of drug was determined as described earlier. Briefly, the POx-Palbo or Palbo-HCl were dispersed in saline (0.1 mg/ml of palbociclib), transferred into floatable Slide-A-Lyzer Mini dialysis device with 3.5 kDa (Thermo Fisher Scientific), and dialyzed against 20 mL PBS containing 10% FBS in compliance with the perfect sink conditions requirements. Four devices were used for each time points. At predetermined time points, the samples were collected and the remaining amount of palbociclib were analyzed by HPLC.

### Animals

*SmoM2^loxP/loxP^* mice were purchased from the Jackson Laboratories (Bar Harbor, ME, USA). *hGFAP-cre* mice were generously provided by Dr. Eva Anton (University of North Carolina, Chapel Hill, NC, USA). *Gli-luc* mice, which express luciferase under the GLI promoter, were generously shared by Dr. Oren Becher and Dr. Eric Holland. Mouse genotyping was performed using *Cre* or *SmoM2* primers. All mice were of the species *Mus musculus* and maintained on a C57BL/6 background over at least 5 generations. Mice were handled under a protocol approved by the University of North Carolina Institutional Animal Care and Use Committee (protocol 19-098).

### Toxicology studies

For MTD studies, we evaluated expected growth of healthy wildtype C57BL/6 mice treated with escalating doses of the drug under study. Drugs were administered by IP injection or oral gavage as indicated to groups of 3 replicate mice, daily from P10-P14 and then every other day until P28. We recorded weights daily and included age-matched littermate controls to determine the expected weight gain over time. MTD was defined as the highest dose resulting in less than 15 % weight gain compared to the control group.

For additional toxicity studies, we treated healthy mice with Palbo-HCl 25 mg/kg or POx-Palbo 25 mg/kg for 15 days (P10~P14: daily, P15~P24: every other day) and compared to saline-injected controls. On P25, mice were sacrificed, and blood were collected and a comprehensive complete blood counts and blood chemistry panel were performed. Major organs including Heart, thymus, lung, liver, spleen and kidney were harvest, fixed in formalin, and subjected to pathological analysis by H&E staining.

### Pharmacokinetic analysis

Groups of 3-4 replicate *G-Smo* mice were administered via intraperitoneal on P10 with a single dose of POx-Palbo or palbociclib-HCl at 25 mg/kg. All samples contained [3H] palbociclib (100 μci/kg). At various time points at 0.083, 2, 4, 8 and 24 hour post injections, three to four *G-Smo* mice from each groups were euthanized, and blood and organs (forebrain, tumor, liver, spleen, and kidney) were collected. Each organs were weighted and homogenized in 1 ml of Soluene-350. 50 μl of serum from each sample was collected and 1 ml of a mixture of soluene-350 and isopropyl alcohol at 1:1 ratio was added. A volume of 15 mL Ultima Gold LLC scintillation cocktail (PerkinElmer, USA) was added to each vial, vortexed, and counted in a liquid scintillation counter, Tricarb 4000 for [3H] palbociclib. Non-compartmental analysis (NCA) was performed using the Phoenix Winonlin software. Parameters derived from the NCA were used as initial estimates in the model building.

### Efficacy studies

*G-Smo* mice were randomized into the indicated treatment groups with 4-12 replicate mice per group and administered via oral or intraperitoneal on P10 with each regimen. The indicated formulations were administered until P35, unless mice first developed symptoms of tumor progression such as hunched posture, ataxia, tremor and seizures.

### Pharmacodynamic analysis

P10-12 tumor mice were administered drugs (POx-Palbo, Palbo-HCl, POx-Sapanisertib, POx-(Palbo+Sapanisertib)) via intraperitoneal injections. EdU (40 mg/kg) was administered to mice thirty minutes before harvest. Tumor tissue was harvested and subjected to cellular dissociation via Worthington Papain Dissociation System kit. Dissociated Tumor cells were fixed for 15 minutes on ice. Washed with FACS wash buffer. Fixed cells were stained with fluorophore markers for DNA (FX Cycle stain, ThermoFisher Scientific, CAT #, F10347), EdU (Click-it Edu Kit, Thermofisher Scientific, CAT# C10337), and phoshpho-RB content (phospho-RB (Ser807/811), Cell Signaling Technology, CAT# 8974S). Stained cells were prepared in appropriate strained for flow analysis.

### Flow Cytometry

Stained cells were resuspended ins Sheath Fluid and ran on a LSR II flow cytometer provided by the UNC Flow Cytometry Core. Proper compensation controls were used.

### Immunofluorescence imaging

Brains including tumors from *G-Smo* pups were harvested, fixed in 4% paraformaldehyde for 48 hours, and embedded in paraffin at the UNC Center for Gastrointestinal Biology and Disease Histology core. Sections were deparaffinized, and antigen retrieval was performed using a low-pH citric acid-based buffer. Staining was performed and stained slides were digitally scanned using the Leica Biosystems Aperio ImageScope software (12.3.3) with assistance from the UNC Translational Pathology Laboratory. The primary antibodies used were anti-pRB (Cell Signaling Technology, CAT # 8516), anti-p4EBP1 (Cell Signaling Technology, CAT #2855), anti-GLT1 (Thermofisher Scientific, CAT #701988).

### Drop-Seq library preparation, sequencing, and analysis

We used Drop-seq (*22*) methods for scRNA sequence study as in our prior studies (*21*, *23*). Briefly, Drop-seq libraries were prepared according to the Drop-seq protocol V 3.1 (*22*) with full details available online (http://mccarrolllab.com/dropseq/). Cell and bead concentrations were both set to between 95 and 110/μL. Tumor cells were co-encapsulated using a Dolomite-brand glass device. All cells were processed within 1 h of tissue dissociation. Flow rates on the glass device were set to 2400 and 12,000 μL/h for cells/beads and oil, respectively, with a 1–2.5% bead occupancy rate. From the obtained library, the raw sequence data were processed in a Linux environment using Drop-seq Tools V1.13 (https://github.com/broadinstitute/Drop-seq/releases) to generate a digital expression (DGE) matrix. DGE matrices were used to generate Seurat objects in R (https://satijalab.org/seurat/). Input data are raw sequences in Fastq format, demultiplexed by sample identity. We first convert Fastq to BAM/SAM format and merge samples that were sequenced across multiple lanes.

The data was normalized using the SCTransform method as implemented in Seurat. PCA was performed on the 3,000 most highly variable using the RunPCA function. We used the FindNeighbors and FindClusters functions to identify cell clusters based on the Louvain algorithm. To identify differential genes between clusters of cells, we used the Wilcoxon rank sum test to compare gene expression of cells within the cluster of interest to all cells outside that cluster, as implemented by the FindMarkers function. Uniform Manifold Approximation and Projection was used to reduce the PCs to two dimensions for data visualization using the RunUMAP function. Biological processes implicated by differential gene expression profiles were identified using GO Enrichment Analysis (*34*–*36*).

## SUPPLEMENTARY MATERIALS

Fig. S1. Scheme of the synthesis of POx block copolymer to be used to prepared POx-micelles.

Fig. S2. Dynamic flow cytometry cell-cycle gating strategy in medulloblastoma-bearing mice.

Fig. S3. (A) Characterization of drug loaded POx micelles. (B-D) Toxicity studies in C57BL/6 mice. The weights of mice treated with (B) Gemcitabine, (C) POx-Etoposide, and (C) POx-Sapanisertib over time. The gray range indicates the mean weights of ± SEM of littermate controls.

Fig. S4. Characterization of (A) POx-(Palbociclib+Vismodegib), (B) POx-(Palbociclib+Etoposide), and (C) POx-(Palbociclib+Sapanisertib) including particle size distribution, zeta potential, and morphology. (D) Particle size, PDI, loading capacity and loading efficiency of two-drug loaded POx micelles.

Table S1. PK parameters of palbociclib in liver, kidney and spleen.

Table S2. Palbociclib regimens used in *in vivo* testing.

Table S3. Palbociclib+vismodegib regimens used in *in vivo* testing.

Table S4. Palbociclib+gemcitabine regimens used in *in vivo* testing.

Table S5. Palbociclib+etoposide regimens used in *in vivo* testing.

Table S6. Palbociclib+Sapanisertib regimens used in *in vivo* testing.

Table S7. HPLC conditions for analyzing the drug concentration in POx micelle.

Data file S1. Gene expression changes and GO analysis.

## Acknowledgments

We thank the UNC CGBID Histology Core supported by P30 DK 034987, the UNC Tissue Pathology Laboratory Core supported by NCI CA016086 and UNC UCRF, and the Chapel Hill Analytical and Nanofabrication Laboratory, supported by the National Science Foundation Grant ECCS-1542015, for help with electron microscopy.

## Author contributions

Conceptualization: CL, TD, TRG, MSP, AVK; Methodology: CL, TD, VLG, DM, JDR, DH; Investigation: CL, TD, VLG, DM; Visualization: CL, TD; Funding acquisition: TRG, MSP; Project administration: TRG, MSP; Supervision: TRG, MSP, AVK; Writing – original draft: CL, TD; Writing – review & editing: TRG, MSP.

## Data and materials availability

All data are available in the main text or the supplementary materials.

