## Supplementary Materials for "Enhancing CDK4/6 inhibitor therapy for medulloblastoma using nanoparticle delivery and scRNA-seq-guided combination with sapanisertib"

Fig. S1. Scheme of the synthesis of POx block copolymer to be used to prepared POx-micelles.

Fig. S2. Dynamic flow cytometry cell-cycle gating strategy in medulloblastoma-bearing mice.

Fig. S3. (A) Characterization of drug loaded POx micelles. (B-D) Toxicity studies in C57BL/6 mice. The weights of mice treated with (B) Gemcitabine, (C) POx-Etoposide, and (C) POx-Sapanisertib over time. The gray range indicates the mean weights of ± SEM of littermate controls.

Fig. S4. Characterization of (A) POx-(Palbociclib+Vismodegib), (B) POx-(Palbociclib+Etoposide), and (C) POx-(Palbociclib+Sapanisertib) including particle size distribution, zeta potential, and morphology. (D) Particle size, PDI, loading capacity and loading efficiency of two-drug loaded POx micelles.

Table S1. PK parameters of palbociclib in liver, kidney and spleen.

Table S2. Palbociclib regimens used in *in vivo* testing.

Table S3. Palbociclib+vismodegib regimens used in *in vivo* testing.

Table S4. Palbociclib+gemcitabine regimens used in *in vivo* testing.

Table S5. Palbociclib+etoposide regimens used in *in vivo* testing.

Table S6. Palbociclib+Sapanisertib regimens used in *in vivo* testing.

Table S7. HPLC conditions for analyzing the drug concentration in POx micelle.

Data file S1. Gene expression changes and GO analysis.

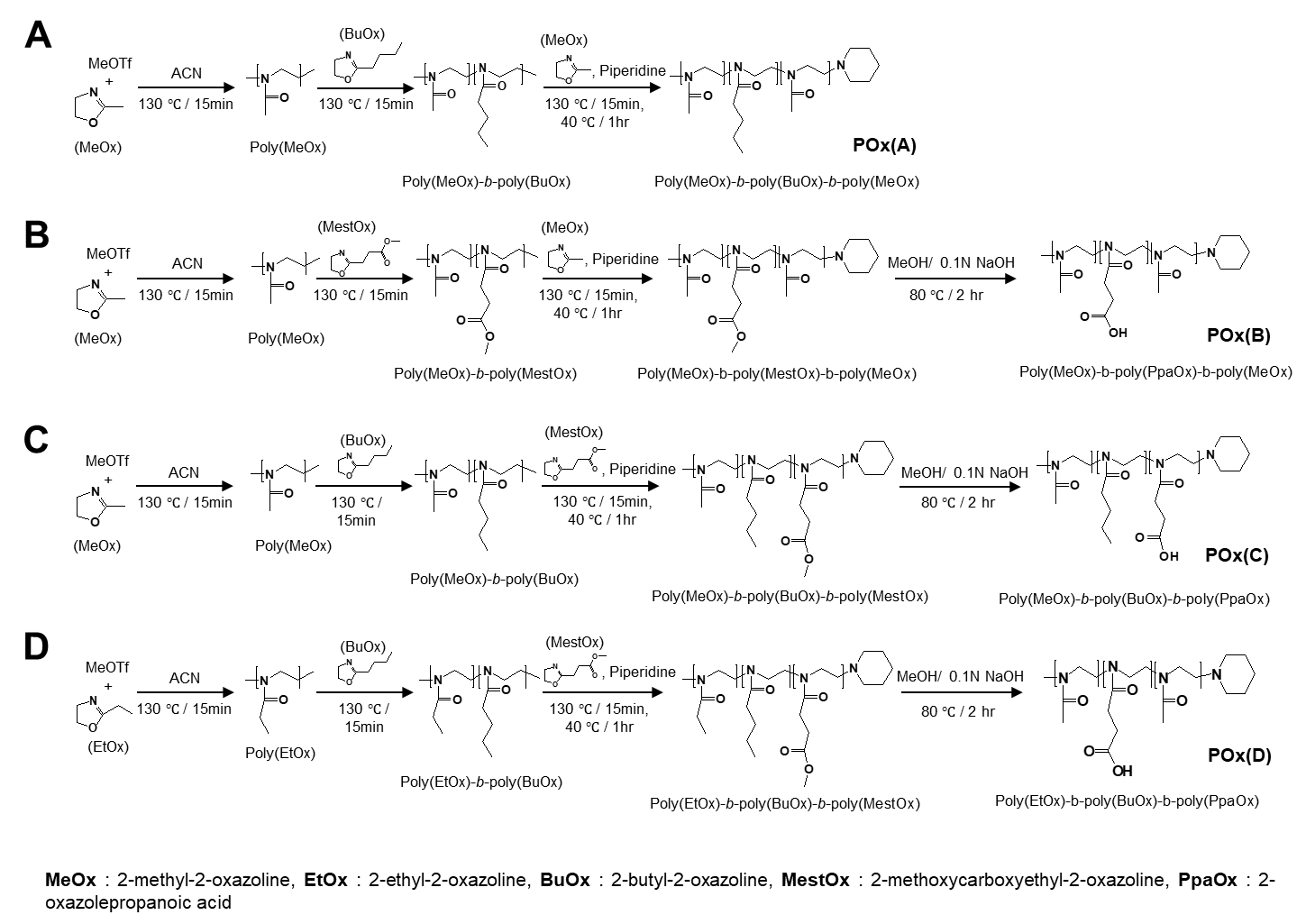

**Fig. S1.** Scheme of the synthesis of various POx block copolymers

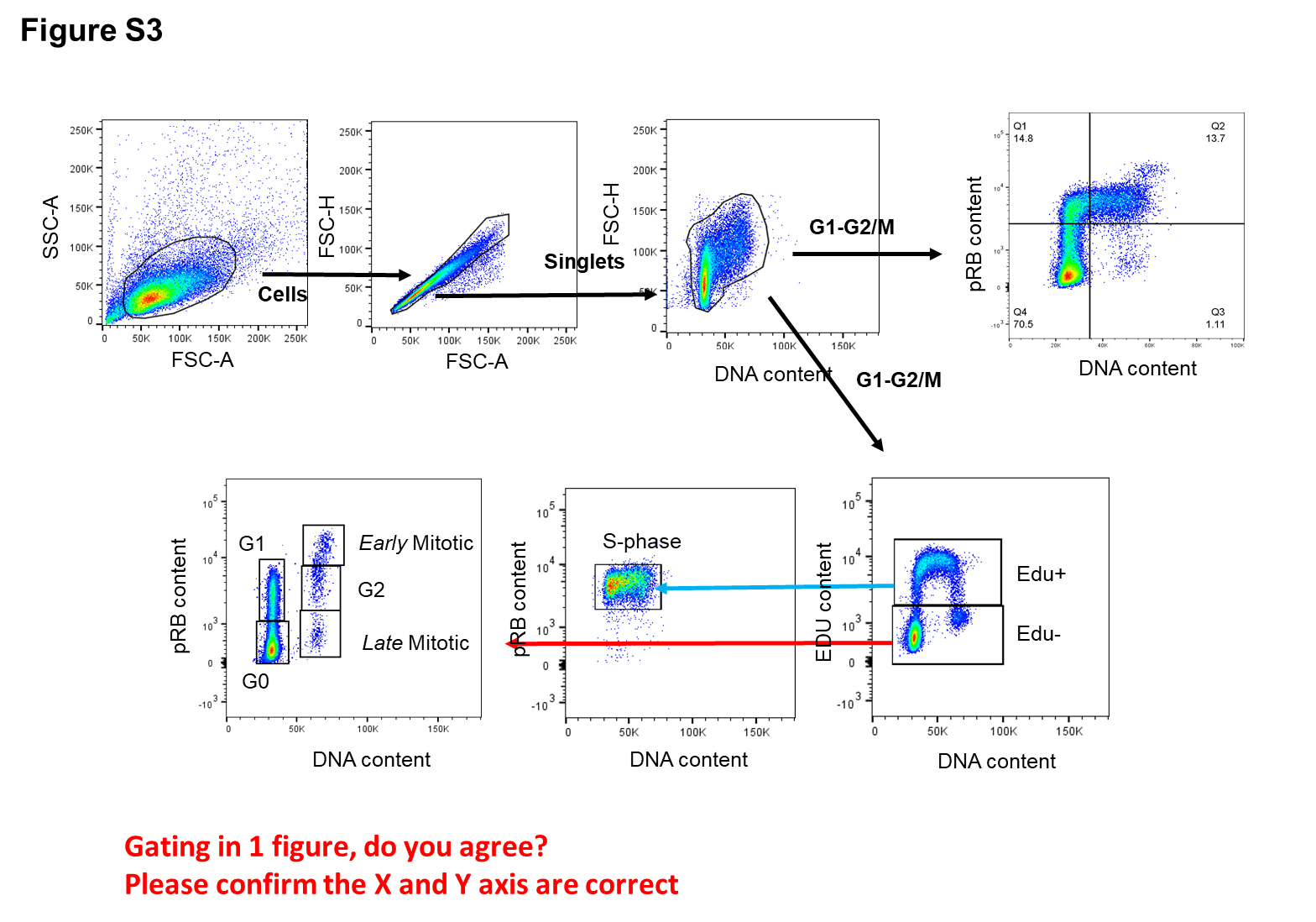

**Fig S2.** Dynamic flow cytometry cell-cycle gating strategy in medulloblastoma-bearing mice.

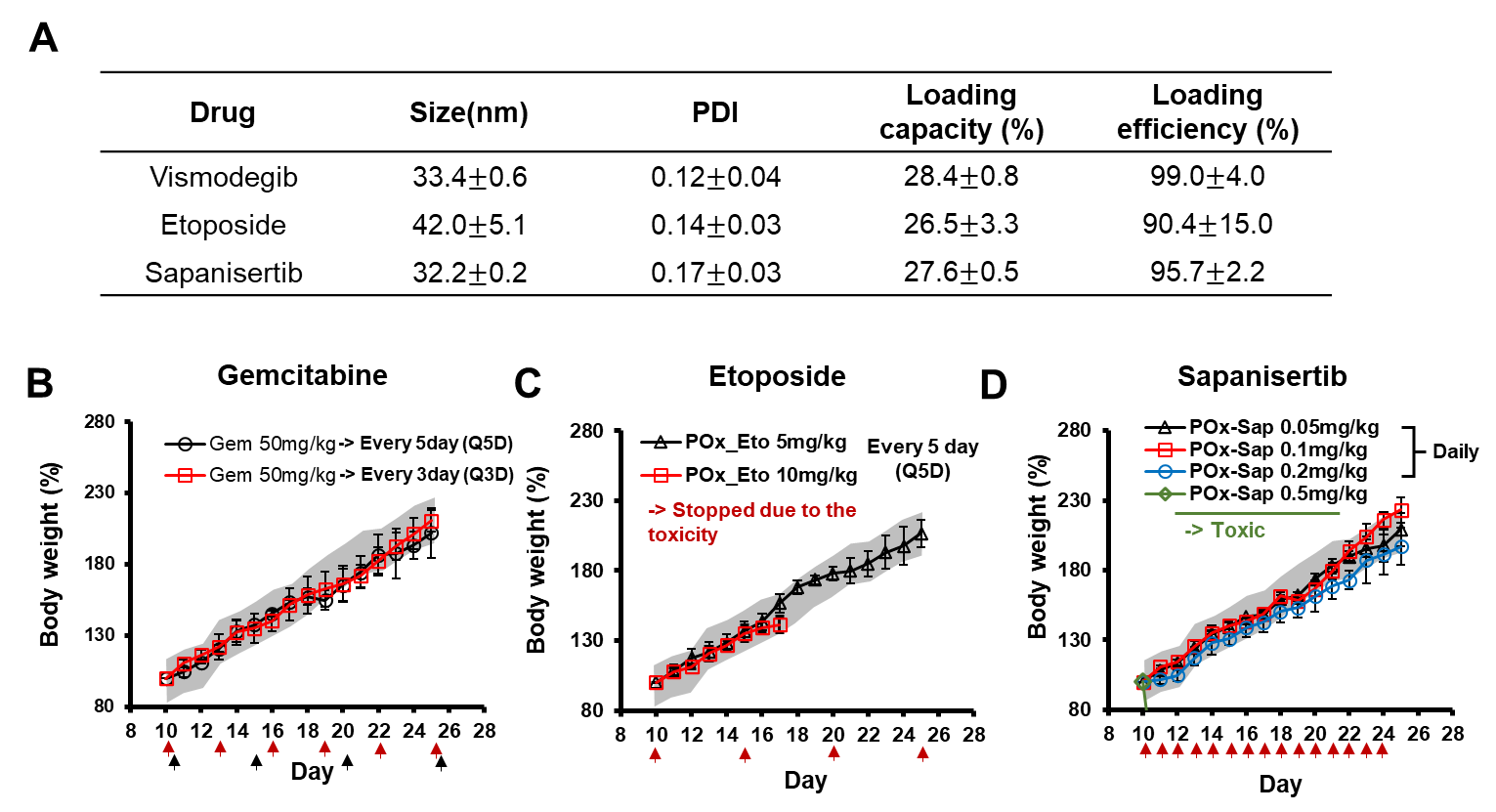

**Fig. S3.** (A) Characterization of drug loaded POx micelles. (B-D) Toxicity studies in C57BL/6 mice. The weights of mice treated with (B) Gemcitabine, (C) POx-Etoposide, and (C) POx-Sapanisertib over time. The gray range indicates the mean weights of $\pm$ SEM of littermate controls.

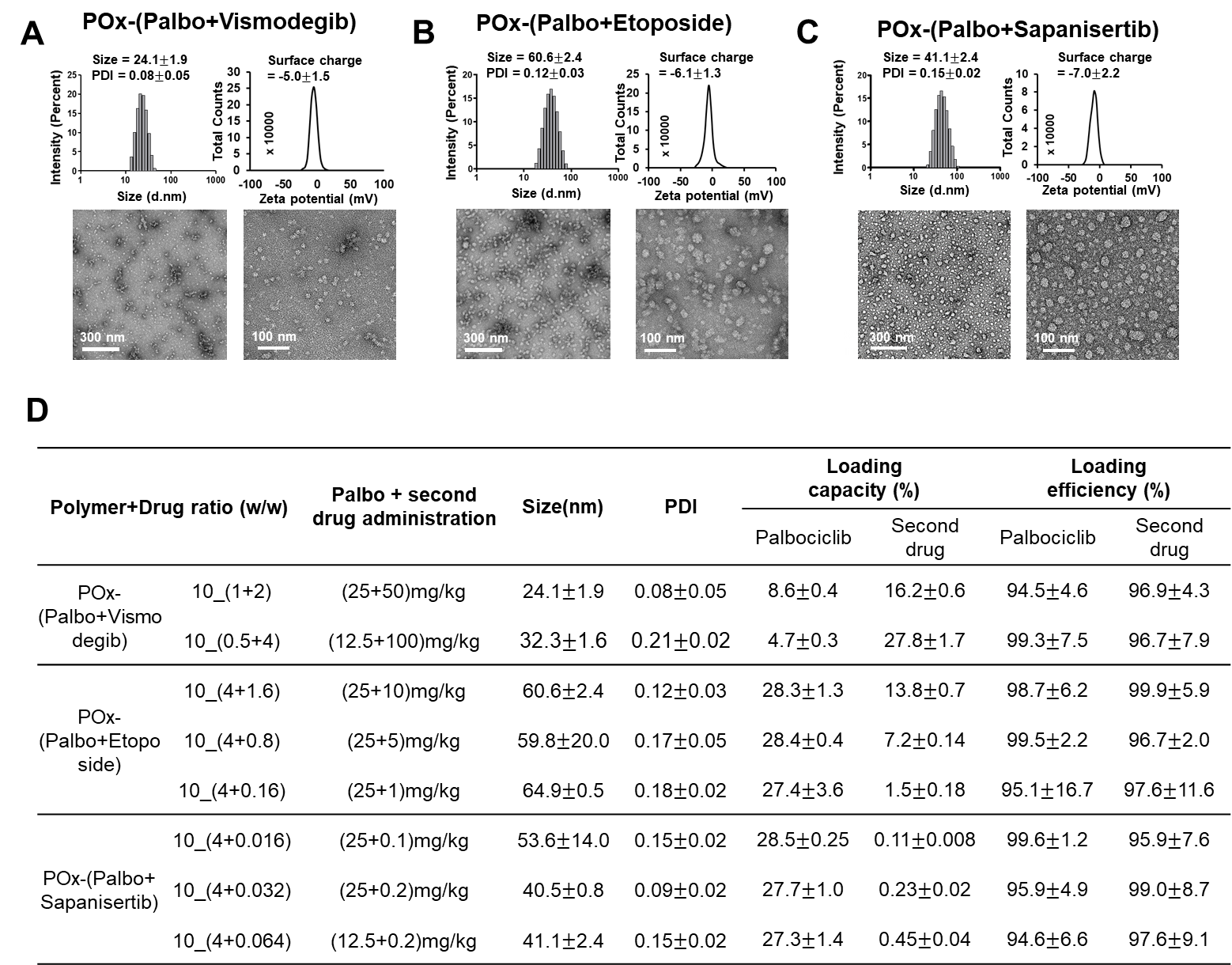

**Fig. S4.** Characterization of (A) POx-(Palbociclib+Vismodegib), (B) POx-(Palbociclib+Etoposide), and (C) POx-(Palbociclib+Sapanisertib) including particle size distribution, zeta potential, and morphology. (D) Particle size, PDI, loading capacity and loading efficiency of two-drug loaded POx micelles.

**Table S1.** PK parametes of given formulations in liver, kidney and spleen. (n = 3 ± SD) *p < 0.05, **p < 0.01, ***p < 0.001.

|  | **POx-Palbo** | | | **Palbo-HCl** | | |
| --- | --- | --- | --- | --- | --- | --- |
|  | **Liver** | **Kidney** | **Spleen** | **Liver** | **Kidney** | **Spleen** |
| **T_max_ (hr) (%ID/ml)** | 4 | 0.08 | 0.08 | 0.08 | 2 | 0.08 |
| **C_max_ (**$\boldsymbol{\mu}$**g/ml)** | 95.68 | 52.71 | 88.26 | 76.34 | 81.85 | 77.30 |
| **AUC_last_ (h**$\boldsymbol{\times\mu}$**g/mL)** | 1109.11 | 766.55 | 318.48 | 682.13 | 632.45 | 305.01 |

**Table S2.** Palbociclib regimens used in *in vivo* testing

| **Day** | **Control** | **Palbociclib-HCl (n=8)** | **POx-Palbociclib** | | | | |
| --- | --- | --- | --- | --- | --- | --- | --- |
|  | **(n=7)** |  | **#1 (n=9)** | **#2 (n=12)** | | | **#3 (n=10)** |
| **10** | No treatment | P.H 10 mg/kg | P 50 mg/kg | P 25 mg/kg | | | P 25 mg/kg |
| **11** |  | P.H 10 mg/kg | P 50 mg/kg | P 25 mg/kg | | | P 25 mg/kg |
| **12** |  | P.H 10 mg/kg | P 50 mg/kg | P 25 mg/kg | | | P 25 mg/kg |
| **13** |  | P.H 10 mg/kg | P 50 mg/kg | P 25 mg/kg | | | P 25 mg/kg |
| **14** |  | P.H 10 mg/kg | P 50 mg/kg | P 25 mg/kg | | | P 12.5 mg/kg |
| **15** |  | X | X | P 25 mg/kg | | | P 12.5 mg/kg |
| **16** |  | P.H 10 mg/kg | P 50 mg/kg | P 25 mg/kg | | | P 12.5 mg/kg |
| **17** |  | X | X | P 25 mg/kg | | | P 12.5 mg/kg |
| **18** |  | P.H 10 mg/kg | P 50 mg/kg | P 25 mg/kg | | | P 12.5 mg/kg |
| **19** |  | X | X | P 25 mg/kg | | | P 12.5 mg/kg |
| **20** |  | P.H 10 mg/kg | P 50 mg/kg | P 25 mg/kg | | | P 12.5 mg/kg |
| **21** |  | X | X | P 25 mg/kg | | | P 12.5 mg/kg |
| **22** |  | P.H 10 mg/kg | P 50 mg/kg | P 25 mg/kg | | | P 12.5 mg/kg |
| **23** |  | X | X | P 25 mg/kg | | | P 12.5 mg/kg |
| **~35** | * | | | | | | |
|  | * | | | | | | |
|  | * | | | | | | |
| **Median** | **16** | **16.5** | **23** | | **22** | **22** | |
| **Survival** |  |  |  |  |  |  |  |

**Table S3.** Palbociclib+vismodegib regimens used in *in vivo* testing

| **Day** | **Palbociclib+Vismodegib (P+V)** | | |
| --- | --- | --- | --- |
|  | **#1 (n=8)** | **#2 (n=8)** | **#3 (n=6)** |
| **10** | P+V (25+50) mg/kg | P+V (25+50) mg/kg | P 25 mg/kg |
| **11** | P+V (25+50) mg/kg | P+V (25+50) mg/kg | P 25 mg/kg |
| **12** | P+V (25+50) mg/kg | P+V (25+50) mg/kg | P 25 mg/kg |
| **13** | P 25 mg/kg | P+V (25+50) mg/kg | P 25 mg/kg |
| **14** | P+V (25+50) mg/kg | P+V (25+50) mg/kg | P+V (12.5+100) mg/kg |
| **15** | P 25 mg/kg | X | P+V (12.5+100) mg/kg |
| **16** | P+V (25+50) mg/kg | P+V (25+50) mg/kg | P+V (12.5+100) mg/kg |
| **17** | P 25 mg/kg | X | X |
| **18** | P+V (25+50) mg/kg | P+V (25+50) mg/kg | P+V (12.5+100) mg/kg |
| **19** | P 25 mg/kg | X | X |
| **20** | P+V (25+50) mg/kg | P+V (25+50) mg/kg | P+V (12.5+100) mg/kg |
| **21** | P 25 mg/kg | X | X |
| **22** | P+V (25+50) mg/kg | P+V (25+50) mg/kg | P+V (12.5+100) mg/kg |
| **23** | P 25 mg/kg | X | X |
| **~35** | * | | |
|  | * | | |
|  | * | | |
| **Median** | **19.5** | **19** | **21** |
| **Survival** |  |  |  |

**Table S4.** Palbociclib+gemcitabine regimens used in *in vivo* testing

| **Day** | **Palbociclib+Gemcitabine (P+G)** | | |
| --- | --- | --- | --- |
|  | **#1 (n=4)** | **#2 (n=7)** | **#3 (n=5)** |
| **10** | P 25 mg/kg | P 25 mg/kg | P 25 mg/kg |
| **11** | P 25 mg/kg | P 25 mg/kg | P 25 mg/kg |
| **12** | P+G (25+50) mg/kg | P 25 mg/kg | P 25 mg/kg |
| **13** | P 25 mg/kg | P 25 mg/kg | P 25 mg/kg |
| **14** | P 25 mg/kg | P+G (25+25) mg/kg | P+G (12.5+25) mg/kg |
| **15** | P+G (25+50) mg/kg | P 25 mg/kg | P 12.5 mg/kg |
| **16** | P 25 mg/kg | P 25 mg/kg | P 12.5 mg/kg |
| **17** | P 25 mg/kg | P 25 mg/kg | P 12.5 mg/kg |
| **18** | P+G (25+50) mg/kg | P 25 mg/kg | P 12.5 mg/kg |
| **19** | P 25 mg/kg | P+G (25+25) mg/kg | P+G (12.5+25) mg/kg |
| **20** | P 25 mg/kg | P 25 mg/kg | P 12.5 mg/kg |
| **21** | P+G (25+50) mg/kg | P 25 mg/kg | P 12.5 mg/kg |
| **22** | P 25 mg/kg | P 25 mg/kg | P 12.5 mg/kg |
| **23** | P 25 mg/kg | P 25 mg/kg | P 12.5 mg/kg |
| **~35** | * | | |
|  | * | | |
|  | * | | |
| **Median** | **16.5** | **20** | **22** |
| **Survival** |  |  |  |

**Table S5.** Palbociclib+etoposide regimens used in *in vivo* testing

|  | **Etoposide** | **Palbociclib+Etoposide (P+E)** | | | | | | | |
| --- | --- | --- | --- | --- | --- | --- | --- | --- | --- |
| **Day** | **#1 (n=10)** | **#1 (n=5)** | **#2 (n=4)** | **#3 (n=13)** | **#4 (n=8)** | **#5 (n=5)** | **#6 (n=8)** | **#7 (n=8)** | **#8 (n=5)** |
| **10** | E 2.5 mg/kg | P 25 mg/kg | P+E (25+2.5) mg/kg | P+E (25+2.5) mg/kg | P 25 mg/kg | P+E (25+10) mg/kg | P+E (25+7.5) mg/kg | P 50 mg/kg | P+E (25+1) mg/kg |
| **11** | X | P 25 mg/kg | P 25 mg/kg | P 25 mg/kg | P 25 mg/kg | P 25 mg/kg | P 25 mg/kg | P 25 mg/kg | P 25 mg/kg |
| **12** | X | P 25 mg/kg | P 25 mg/kg | P 25 mg/kg | P 25 mg/kg | P 25 mg/kg | P 25 mg/kg | P 25 mg/kg | P 25 mg/kg |
| **13** | X | P 25 mg/kg | P 25 mg/kg | P 25 mg/kg | P 25 mg/kg | P 25 mg/kg | P 25 mg/kg | P+E (25+5) mg/kg | P+E (25+1) mg/kg |
| **14** | X | P+E (25+5) mg/kg | P 25 mg/kg | P 25 mg/kg | P+E (25+2.5) mg/kg | P 25 mg/kg | P 25 mg/kg | P 25 mg/kg | P 25 mg/kg |
| **15** | E 2.5 mg/kg | P 25 mg/kg | P+E (12.5+2.5) mg/kg | P+E (25+2.5) mg/kg | P 25 mg/kg | P+E (25+10) mg/kg | P+E (25+7.5) mg/kg | X | P 25 mg/kg |
| **16** | X | P 25 mg/kg | P 12.5 mg/kg | P 25 mg/kg | P 25 mg/kg | P 25 mg/kg | P 25 mg/kg | X | P+E (25+1) mg/kg |
| **17** | X | P 25 mg/kg | P 12.5 mg/kg | P 25 mg/kg | P 25 mg/kg | P 25 mg/kg | P 25 mg/kg | P 25 mg/kg | P 25 mg/kg |
| **18** | X | P 25 mg/kg | P 12.5 mg/kg | P 25 mg/kg | P 25 mg/kg | P 25 mg/kg | P 25 mg/kg | P 25 mg/kg | P 25 mg/kg |
| **19** | X | P+E (25+5) mg/kg | P 12.5 mg/kg | P 25 mg/kg | P+E (25+2.5) mg/kg | P 25 mg/kg | P 25 mg/kg | P 25 mg/kg | P+E (25+1) mg/kg |
| **20** | E 2.5 mg/kg | P 25 mg/kg | P+E (12.5+2.5) mg/kg | P+E (25+2.5) mg/kg | P 25 mg/kg | P+E (25+10) mg/kg | P+E (25+7.5) mg/kg | P+E (25+5) mg/kg | P 25 mg/kg |
| **21** | X | P 25 mg/kg | P 12.5 mg/kg | P 25 mg/kg | P 25 mg/kg | P 25 mg/kg | P 25 mg/kg | P 25 mg/kg | P 25 mg/kg |
| **22** | X | P 25 mg/kg | P 12.5 mg/kg | P 25 mg/kg | P 25 mg/kg | P 25 mg/kg | P 25 mg/kg | X | P+E (25+1) mg/kg |
| **23** | X | P 25 mg/kg | P 12.5 mg/kg | P 25 mg/kg | P 25 mg/kg | P 25 mg/kg | P 25 mg/kg | X | P 25 mg/kg |
| **~35** | *  *  * | | | | | | | | |
| **Median**  **Survival** | **19.5** | **20** | **19** | **24** | **21** | **18** | **18.5** | **18** | **24** |

**Table S6.** Palbociclib+Sapanisertib regimens used in *in vivo* testing

|  | **Sapanisertib** | **Palbociclib+Sapanisertib (P+S)** | | | |
| --- | --- | --- | --- | --- | --- |
| **Day** | **#1 (n=9)** | **#2 (n=5)** | **#1 (n=4)** | **#2 (n=10)** | **#3 (n=10)** |
| **10** | S 0.1 mg/kg | S 0.2 mg/kg | P+S (25+0.1) mg/kg | P 25 mg/kg | P 25 mg/kg |
| **11** | S 0.1 mg/kg | S 0.2 mg/kg | P+S (25+0.1) mg/kg | P 25 mg/kg | P 25 mg/kg |
| **12** | S 0.1 mg/kg | S 0.2 mg/kg | P+S (25+0.1) mg/kg | P 25 mg/kg | P 25 mg/kg |
| **13** | S 0.1 mg/kg | S 0.2 mg/kg | P+S (25+0.1) mg/kg | P 25 mg/kg | P 25 mg/kg |
| **14** | S 0.1 mg/kg | S 0.2 mg/kg | P+S (25+0.1) mg/kg | P+S (12.5mg+0.1) mg/kg | P+S (12.5mg+0.2) mg/kg |
| **15** | S 0.1 mg/kg | S 0.2 mg/kg | P+S (25+0.1) mg/kg | P+S (12.5mg+0.1) mg/kg | P+S (12.5mg+0.2) mg/kg |
| **16** | S 0.1 mg/kg | S 0.2 mg/kg | P+S (25+0.1) mg/kg | P+S (12.5mg+0.1) mg/kg | P+S (12.5mg+0.2) mg/kg |
| **17** | S 0.1 mg/kg | S 0.2 mg/kg | P+S (25+0.1) mg/kg | P+S (12.5mg+0.1) mg/kg | X |
| **18** | S 0.1 mg/kg | S 0.2 mg/kg | P+S (25+0.1) mg/kg | P+S (12.5mg+0.1) mg/kg | P+S (12.5mg+0.2) mg/kg |
| **19** | S 0.1 mg/kg | S 0.2 mg/kg | P+S (25+0.1) mg/kg | P+S (12.5mg+0.1) mg/kg | X |
| **20** | S 0.1 mg/kg | S 0.2 mg/kg | P+S (25+0.1) mg/kg | P+S (12.5mg+0.1) mg/kg | P+S (12.5mg+0.2) mg/kg |
| **21** | S 0.1 mg/kg | S 0.2 mg/kg | P+S (25+0.1) mg/kg | P+S (12.5mg+0.1) mg/kg | X |
| **22** | S 0.1 mg/kg | S 0.2 mg/kg | P+S (25+0.1) mg/kg | P+S (12.5mg+0.1) mg/kg | P+S (12.5mg+0.2) mg/kg |
| **23** | S 0.1 mg/kg | S 0.2 mg/kg | P+S (25+0.1) mg/kg | P+S (12.5mg+0.1) mg/kg | X |
| **~35** | * | | | | |
|  | * | | | | |
|  | * | | | | |
| **Median** | **21** | **17** | **19.5** | **24** | **> 35 days** |
| **Survival** |  |  |  |  |  |

**Table S7.** HPLC conditions for analyzing the drug concentration in POx micelle

| **Sample** | **Retention time** | | **Detection wavelength (nm)** | **Mobile phase  (Mixture of acetonitrile/water, v/v)** |
| --- | --- | --- | --- | --- |
|  | **Palbociclib** | **Combined drug** |  |  |
| POx-(Palbo+Vismodegib) | 3.2 min | 4.4 min | 280 / 280 | 50%:50% with 0.01% TFA |
| POx-(Palbo+Etoposide) | 3.2 min | 3.8 min | 280 / 227 | 50%:50% with 0.01% TFA |
| POx-(Palbo+Sapanisertib) | 4.1 min | 3.5 min | 280 / 280 | 40%:60% with 0.01% TFA |
